# Adaptive subsets limit the anti-tumoral NK-cell activity in Hepatocellular Carcinoma

**DOI:** 10.1101/2021.04.16.440140

**Authors:** Charlotte Rennert, Catrin Tauber, Pia Fehrenbach, Kathrin Heim, Dominik Bettinger, Oezlem Sogukpinar, Anita Schuch, Britta Franziska Zecher, Bertram Bengsch, Sven A. Lang, Peter Bronsert, Niklas K. Björkström, Stefan Fichtner-Feigl, Michael Schultheiss, Robert Thimme, Maike Hofmann

**Affiliations:** Department of Medicine II, University Hospital Freiburg - Faculty of Medicine, University of Freiburg, Hugstetter Straße 55, Freiburg 79106, Germany; Faculty of Biology, University of Freiburg, Schänzlestraße 1, Freiburg 79104, Germany; I. Department of Medicine, University Medical Centre Hamburg-Eppendorf, Martinistraße 9 52, 20246 Hamburg, Germany; Department of General and Visceral Surgery, University Hospital Freiburg – Faculty of Medicine, Medical Center, University of Freiburg, Hugstetter Straße 55, Freiburg 79106, Germany; Department of General, Visceral and Transplantation Surgery, University Hospital Aachen – Faculty of Medicine, Medical Center, University of Aachen, Pauwelsstraße 30, Aachen 52074, Germany; Institute of Pathology, University Hospital Freiburg - Faculty of Medicine, University of Freiburg, Hugstetter Straße 55, Freiburg 79106, Germany; Tumorbank Comprehensive Cancer Center Freiburg, Medical Center – Faculty of Medicine, University of Freiburg, Hugstetter Straße 55, Freiburg 79106, Germany; Center for Infectious Medicine, Department of Medicine Huddinge, Karolinska Institutet, Karolinska University Hospital, Alfred Nobels allé 8, 141 52 Huddinge, Sweden

**Keywords:** HCC, liver cirrhosis, HCMV, HBV

## Abstract

**Background and Aims:** Hepatocellular carcinoma (HCC) is a global health burden with increasing incidence, poor prognosis and limited therapeutic options. Natural killer (NK) cells exhibit potent anti-tumoral activity and therefore represent potential targets for immunotherapeutic approaches in HCC treatment. However, the human NK-cell repertoire is highly diverse including conventional and adaptive NK cells that differ in phenotype and effector function. Adaptive NK-cell frequencies are increased in association with HCMV (human cytomegalovirus) seropositivity that is also common in HCC patients. In this study, we aimed to gain a better understanding of the NK-cell repertoire and the associated anti-tumoral activity in HCC patients.

**Methods:** In-depth phenotypic and functional flow-cytometry analyses of the HCMV-associated NK cell-repertoire obtained from 57 HCC patients, 33 liver cirrhosis patients and 36 healthy donors (HD).

**Results:** First, adaptive subsets are present in all three cohorts with conserved characteristics in patients with liver cirrhosis and HCC. Second, adaptive NK cells can be isolated from HCC tissue however lack features of tissue-residency and thus probably represent circulating/infiltrating lymphocytes. Third, the anti-tumoral activity by adaptive NK cells is reduced compared to conventional NK-cell subsets, also in HCC. Lastly, frequencies of adaptive NK cells were increased in patients suffering from Hepatitis B virus-associated HCC providing a link between etiology and the NK-cell repertoire in HCC.

**Conclusion:** Adaptive NK cells limit the anti-tumoral activitity of NK cells in HCC, especially in association with HBV infection that is accompanied by an expansion of this NK cell subset.

**Lay summary:** In patients with liver cancer (HCC), a subset of natural killer cells, so called adaptive NK cells show a diminished anti-tumoral activity compared to other, called conventional NK cells. Adaptive NK cells are expanded in patients with HCC associated to Hepatitis B virus infection. Thus, presence of adaptive NK cells contributes to the immune escape in HCC.

## Introduction

Hepatocellular carcinoma (HCC) is the most common form of primary liver cancer in adults and represents a major health problem due to the increasing incidence worldwide [1, 2]. The development of liver cancer is a multifactorial process in which HCC develops from chronic liver damage resulting in pre-malignant cirrhotic remodeling. Chronic liver damage is frequently caused by alcohol consumption, the metabolic syndrome and viral infections [3]. Especially, chronic hepatitis B virus (HBV) infection is the world’s leading cause of HCC, whereas chronic hepatitis C virus (HCV) infection is currently the dominant reason for HCC in the western world [3]. Given the poor prognosis and limited therapeutic options [1], there is an urgent need for new approaches in the treatment of HCC. Several studies have indicated a role of innate and adaptive immunity in HCC progression and control [2-4] and therefore there is growing interest to treat HCC patients with immunotherapeutic approaches. In particular, a longer progression-free survival of HCC patients has been associated with the presence of CD8^+^ T cells targeting tumor-associated antigens [5] and increased NK-cell frequencies in the periphery and within the tumor [6, 7]. These data support that both CD8^+^ T cells and NK cells are potent effector cells of the anti-tumoral immune response and render these immune cells promising targets for treating HCC.

Unlike CD8^+^ T cells, tumor surveillance by NK cells is not mediated by targeting tumor-associated antigens and neo-antigens that are heterogeneously expressed and largely unknown in cancer patients. NK cells mediate anti-tumoral immunity by sensing the down-regulated MHC class I expression [8] (“missing-self response”) [9] and increased expression of cell stress ligands (“altered-self response”, e.g. DNA-damage response induced NKG2D ligands on tumor cells [10, 11]). Furthermore, NK cells are activated in response to cytokines [12-14] present in the tumor microenvironment and CD16 expressed on NK cells, mediates antibody-dependent cellular cytotoxicity (ADCC) [15]. Due to these modes of activation, NK cells represent broadly acting effector cells in tumor surveillance [16].

Generally, the human liver is enriched for NK cells that account for up to 40% of all liver immune cells [17-19] and similarly high frequencies of NK cells can be found in human liver tumors including HCC [20]. In HCC, phenotypic alterations within the NK-cell repertoire as well as diminished NK-cell effector functions have been reported [21], indicating that failure of the NK-cell response may contribute to tumor growth. Furthermore, distinct phenotypic NK-cell characteristics, e.g. high expression of cytotoxic granules, have been associated with a better survival after curative treatment [22]. However, a detailed understanding of the NK-cell response in the context of HCC is missing, especially in light of the emerging knowledge about NK-cell heterogeneity [23].

NK cells represent a heterogeneous population of innate lymphocytes with phenotypically and functionally distinct subsets [24]. In peripheral blood, CD56^dim^ NK cells represent the predominant NK-cell population [25], characterized by a high expression of CD16 and only an intermediate level of CD56. They produce cytokines and have a cytotoxic effect [26]. CD56^bright^ NK cells produce cytokines to a high degree, but do not possess cytotoxic properties [26]. The proportion of CD56^bright^ NK cells in the human liver is significantly higher than in peripheral blood and they are largely composed of liver-resident subsets [27-29]. These liver-resident NK cells exhibit tolerogenic features that partially account for an impaired anti-tumoral NK-cell response in HCC [20, 28]. In contrast, the impact of different CD56^dim^ NK-cell subsets on the anti-tumoral activity in solid tumors like HCC is still unknown.

Diversification of CD56^dim^ NK cells is best described in association with human cytomegalovirus (HCMV) infection resulting in the emergence of adaptive NK cells that are characterized by a distinct epigenetic, phenotypic and functional profile, including downregulation of the signaling adapter FcεRIγ, a decreased cytokine responsiveness but higher ADCC compared to conventional CD56^dim^ NK cells [30-34]. Previously, our group showed that the emergence of HCMV-associated adaptive NK cells shapes the overall NK-cell response in HBV patients [35]. In this study, we now show that adaptive NK cells are present in the peripheral blood of HCMV^+^ HCC patients with liver cirrhosis and HD with conserved characteristics. Furthermore, adaptive NK cells are also detectable within HCC tumor tissue exhibiting a similar profile as in the blood and lacking characteristics of tissue-residency. Hence, adaptive NK cells most probably represent circulating/infiltrating lymphocytes. Compared to conventional NK cells, the anti-tumoral activity of adaptive NK cells is impaired. In line with our previous data, adaptive NK cells are expanded in association with HBV infection also in the context of HCC linking the NK-cell repertoire to HCC etiology.

Taken together, these results indicate that adaptive CD56^dim^ NK cells may contribute to the impaired anti-tumoral NK-cell activity in HCC, especially in association with HBV infection.

## Results

### CD56^dim/bright^ subset diversification of circulating NK cells is maintained in HCC

To assess the CD56^dim/bright^ NK-cell subset distribution in HCC versus patients with liver cirrhosis and HD, we compared the frequencies of circulating CD56^dim^ and CD56^bright^ NK cells (Fig. 1A) in these cohorts. As shown in Fig. 1B/C, CD56^dim^ and CD56^bright^ NK-cell frequencies within the circulating NK-cell population (defined as CD3^-^CD56^+^) were similar in patients with HCC compared to controls (irrespective of HCMV serostatus) with a clear dominance of CD56^dim^ NK cells. However, significantly less CD56^dim^ NK cells were detectable in cirrhotic patients compared to HD (Fig 1D). Furthermore, frequencies of CD56^dim^ and CD56^bright^ NK-cell populations were also not affected by HCMV infection (Fig. 1D/E), a main driver of NK-cell repertoire formation and present in the majority of our HCC cohort (Fig. 1F). Thus, CD56^dim/bright^ subset diversification of circulating NK cells is maintained in HCC.

**Fig. 1.**
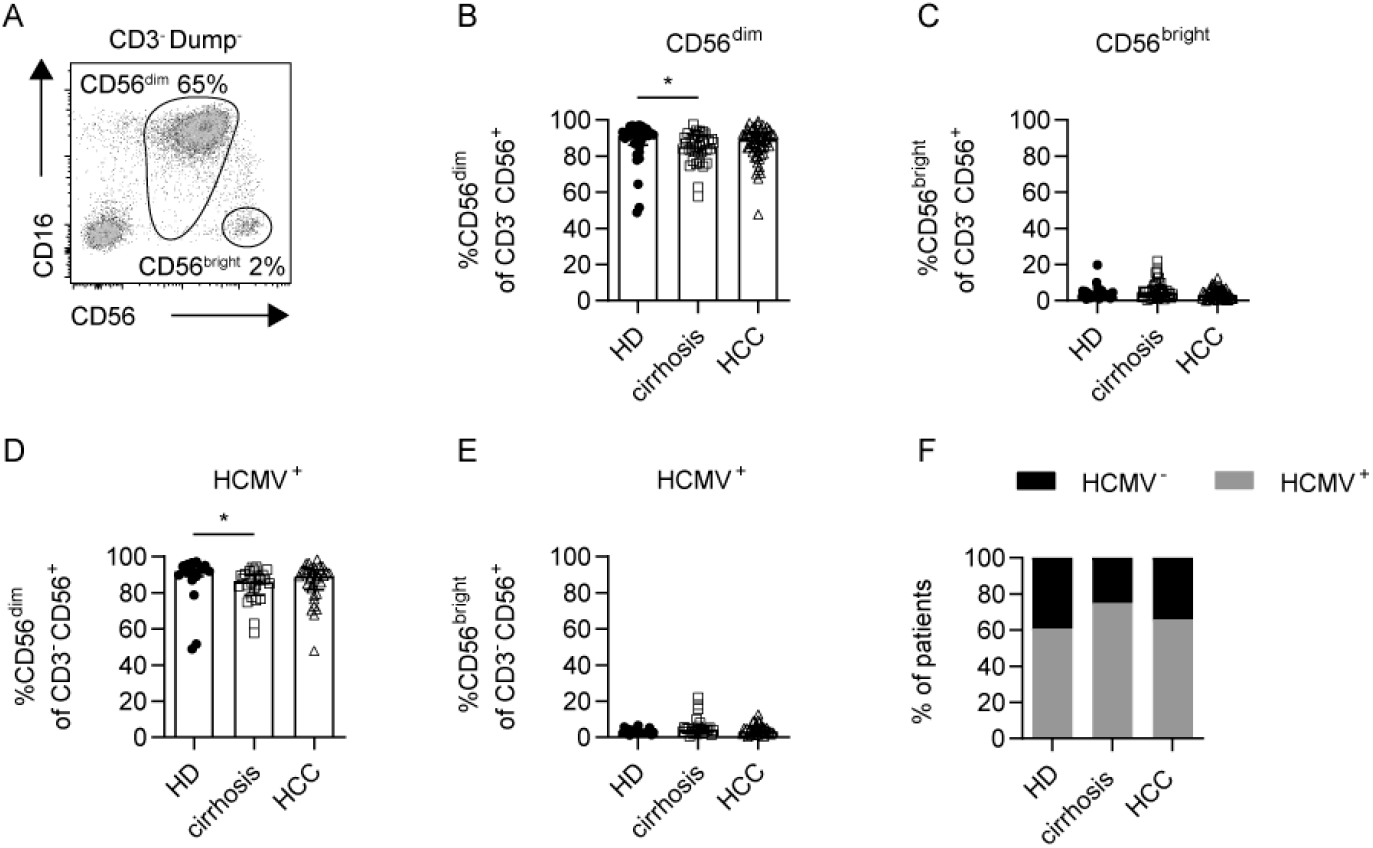
Similar frequency of CD56^dim^ and CD56^bright^ NK cells in peripheral blood of HCC patients compared to controls. Gating strategy (A). Frequency of CD56^dim^ (B) and CD56^bright^ (C) NK cells of CD3^-^CD56^+^ lymphocytes in the peripheral blood of HCC patients, HD and patients with liver cirrhosis. Frequency of CD56^dim^ (D) and CD56^bright^ (E) NK cells in HCMV^+^ individuals. Percentage of HCMV^+^ (grey) and HCMV^-^(black) patients (F). Each dot represents an individual. Bars indicate the median with IQR. Statistical significance was assessed by using Kruskal-Wallis test (B-E). HCMV: human cytomegalovirus, HCC: patients with hepatocellular carcinoma, HD: healthy donors, cirrhosis: patients with liver cirrhosis. Dump^-^cells include doublets, dead cells, CD14^+^ and CD19^+^ cells.

### Profiles of adaptive NK cells are conserved in HCC

Since HCMV seropositivity is frequent, we next wanted to gain a deeper understanding of HCMV-driven CD56^dim^ NK-cell heterogeneity in HCMV^+^ HCC patients. For this, we analyzed expression levels of molecules that have been associated with the HCMV-induced emergence of adaptive NK cells. Specifically, we analyzed the signaling adapters FcεRIγ and Syk, the transcription factors PLZF and Helios, and expression of the surface molecules NKG2C and CD57 [30-34, 36] in HCMV^+^ individuals. Of note, we could detect FcεRIγ^-^, Syk^-^, PLZF^-^, Helios^-^, NKG2C^+^ and CD57^+^ (Fig. 2A) cells within CD56^dim^ NK-cell populations in HCMV^+^ HCC patients. Thus, HCMV-driven CD56^dim^ NK-cell diversification is also evident in HCMV^+^ HCC patients. Importantly, there was no significant difference in the frequencies of the respective NK-cell population in HCMV^+^ HCC patients compared to HCMV^+^ patients with liver cirrhosis and HCMV^+^ HD (Fig. 2A marked in grey).

**Fig. 2:**
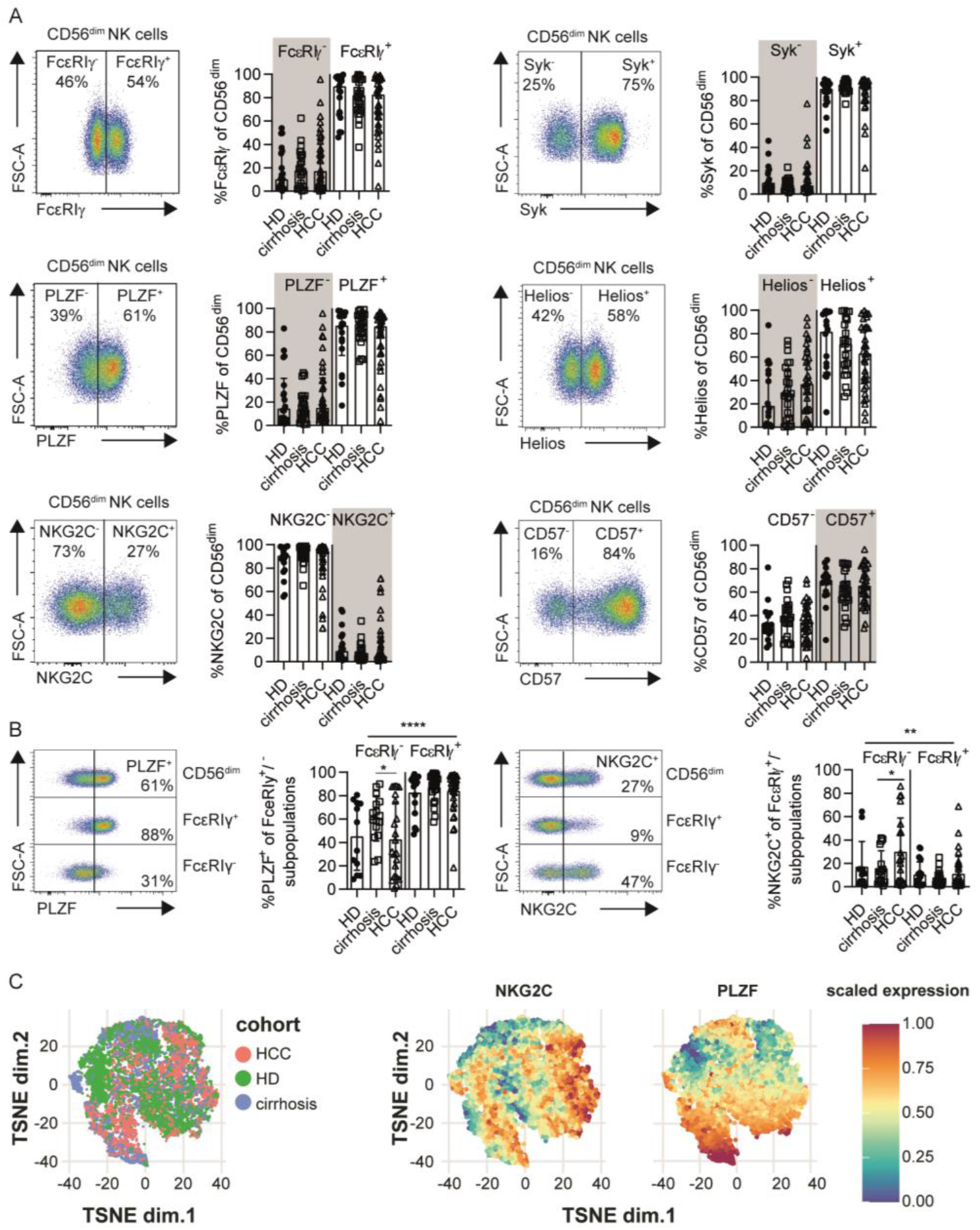
Adaptive NK cell signatures are conserved in HCC patients. FcεRIγ, Syk, PLZF, Helios, NKG2C and CD57 expression on CD56^dim^ NK cells in HCMV^+^ HCC patients (A). PLZF and NKG2C expression of FcεRIγ^-^and FcεRIγ^+^ subpopulations in HCC patients, HD and patients with liver cirrhosis (B). t-SNE representation analysis of concatenated flow cytometry data obtained from FcεRIγ^-^CD56^dim^ NK cells in HCC patients, HD and patients with liver cirrhosis. Expression levels of NKG2C and PLZF are plotted on the t-SNE plots (C). Cells in the grey box correspond to the phenotype of adaptive NK cells. Each dot represents an HCMV^+^ individual. Bars indicate median with IQR. Statistical significance for cohort comparison was assessed by using Kruskal-Wallis test (A) and for subset comparison two-way ANOVA (B). Patients with less than 10% adaptive FcεRIγ^-^CD56^dim^ NK cells were excluded (B). HCC: patients with hepatocellular carcinoma, HD: healthy donors, cirrhosis: patients with liver cirrhosis.

Next, we assessed if the combined phenotype of adaptive NK cells is conserved in HCMV^+^ HCC patients. The presence of adaptive CD56^dim^ NK cells was defined by more than 10% FcεRIγ^-^cells among CD56^dim^ NK-cell populations. For this, we comparatively analyzed the expression of PLZF and NKG2C (Fig. 2B) in FcεRIγ^-^CD56^dim^ NK cells that are significantly more frequent in HCMV^+^ compared to HCMV^-^HCC patients (SI Fig. 1). As previously described [30, 31], lower frequencies of adaptive FcεRIγ^-^CD56^dim^ NK cells compared to conventional FcεRIγ^+^ CD56^dim^ NK cells expressed PLZF (Fig. 2B) whereas higher frequencies expressed NKG2C (Fig. 2B). Importantly, the expression pattern was similar in HCMV^+^ HCC patients compared to HCMV^+^ control cohorts. T-SNE analysis of adaptive FcεRIγ^-^CD56^dim^ NK cells revealed, that the majority of the cells intermingle between all three cohorts (Fig. 2C). Thus, HCMV-driven adaptive NK-cell diversification is conserved in HCMV^+^ HCC compared to HCMV^+^ patients with liver cirrhosis or HCMV^+^ HD. Further phenotypic analysis of CD2, CD7, Siglec-7, CX3CR1 and CXCR3 expression (SI Fig 2) also showed a conserved adaptive NK cell profile in HCMV^+^ HCC patients.

### Adaptive NK cells are present in tumor and non-tumor liver tissue

Human liver is enriched for NK cells that exhibit a considerably different profile compared to circulating NK cells [29, 37]. Thus, we analyzed adjacent non-tumor liver tissue and HCC tumor samples for the presence of adaptive FcεRIγ^-^CD56^dim^ NK cells in HCMV^+^ patients. For this, we stained for FcεRIγ (Fig. 3A), PLZF (Fig. 3B), Helios (Fig. 3C), NKG2C (Fig. 3D) and CD57 (Fig. 3E) in CD56^dim^ NK cells obtained from liver tissues. Indeed, we could detect CD56^dim^ NK cells lacking FcεRIγ, PLZF or Helios and expressing NKG2C or CD57 in adjacent non-tumoral liver tissue and HCC tumor tissue, clearly showing that adaptive NK cells are not restricted to the circulation. To assess whether adaptive NK cells isolated from the tumor are phenotypically different, we compared the expression of PLZF, Helios, NKG2C and CD57 in FcεRIγ^-^CD56^dim^ NK cells from matched blood, adjacent non-tumor liver and HCC tumor tissues obtained from seven HCMV^+^ HCC patients with more than 10% FcεRIγ^-^adaptive CD56^dim^ NK cells in the blood (Fig. 4A). All analyzed markers were equally expressed in adaptive NK cells from the three compartments, suggesting conserved profiles even within HCC tumors. This finding was further corroborated by t-SNE analysis of FcεRIγ^-^adaptive CD56^dim^ NK, showing no clear separation between the cells of the different compartments (Fig. 4B). To further address whether FcεRIγ^-^CD56^dim^ NK cells are tissue/tumor-infiltrating cells or rather represent tissue-resident NK cells, we analyzed the expression of the tissue-resident marker molecules CD69, CXCR6 and CD49a on CD56^bright^ NK cells and CD56^dim^ NK-cell subpopulations obtained from adjacent non-tumor and tumor lesions of HCC patients (Fig. 4C and SI Fig. 3). FcεRIγ^-^adaptive CD56^dim^ NK cells, isolated from liver tissue irrespective of non-tumor or tumor origin, displayed low expression of these tissue-resident markers. Hence, adaptive CD56^dim^ NK cells found in HCC lesions are most probably circulating tumor-infiltrating lymphocytes.

**Fig. 3:**
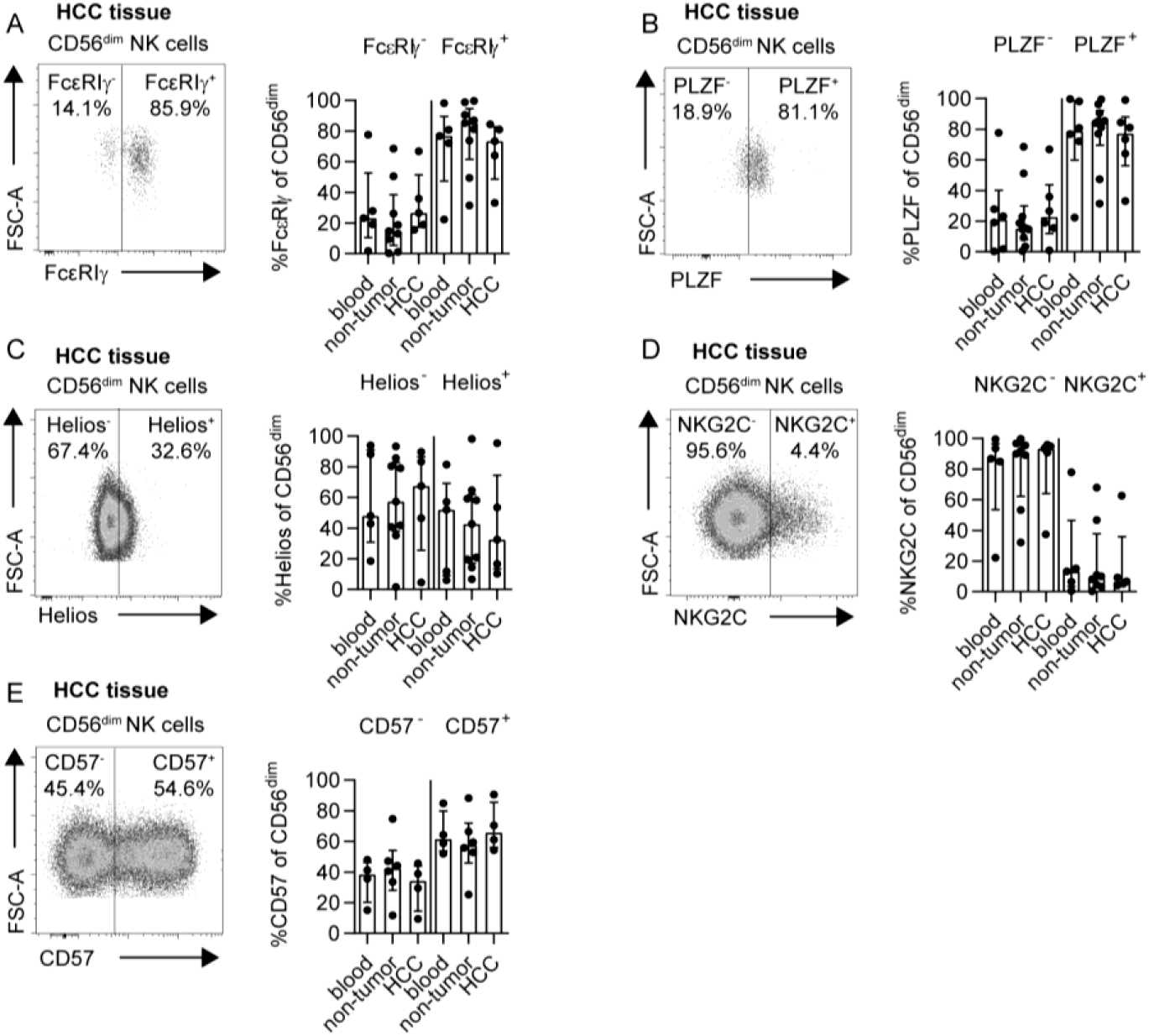
NK cell repertoire is conserved in blood and tissue in HCC patients. FcεRIγ (A), PLZF (B), Helios (C), NKG2C (D) and CD57 (E) expression on CD56^dim^ NK cells in HCC tissue, adjacent non-tumoral liver tissue and matched blood samples. Bars indicate median with IQR. Statistical significance for cohort comparison was tested using Kruskal-Wallis test. HCC: hepatocellular carcinoma.

**Fig. 4:**
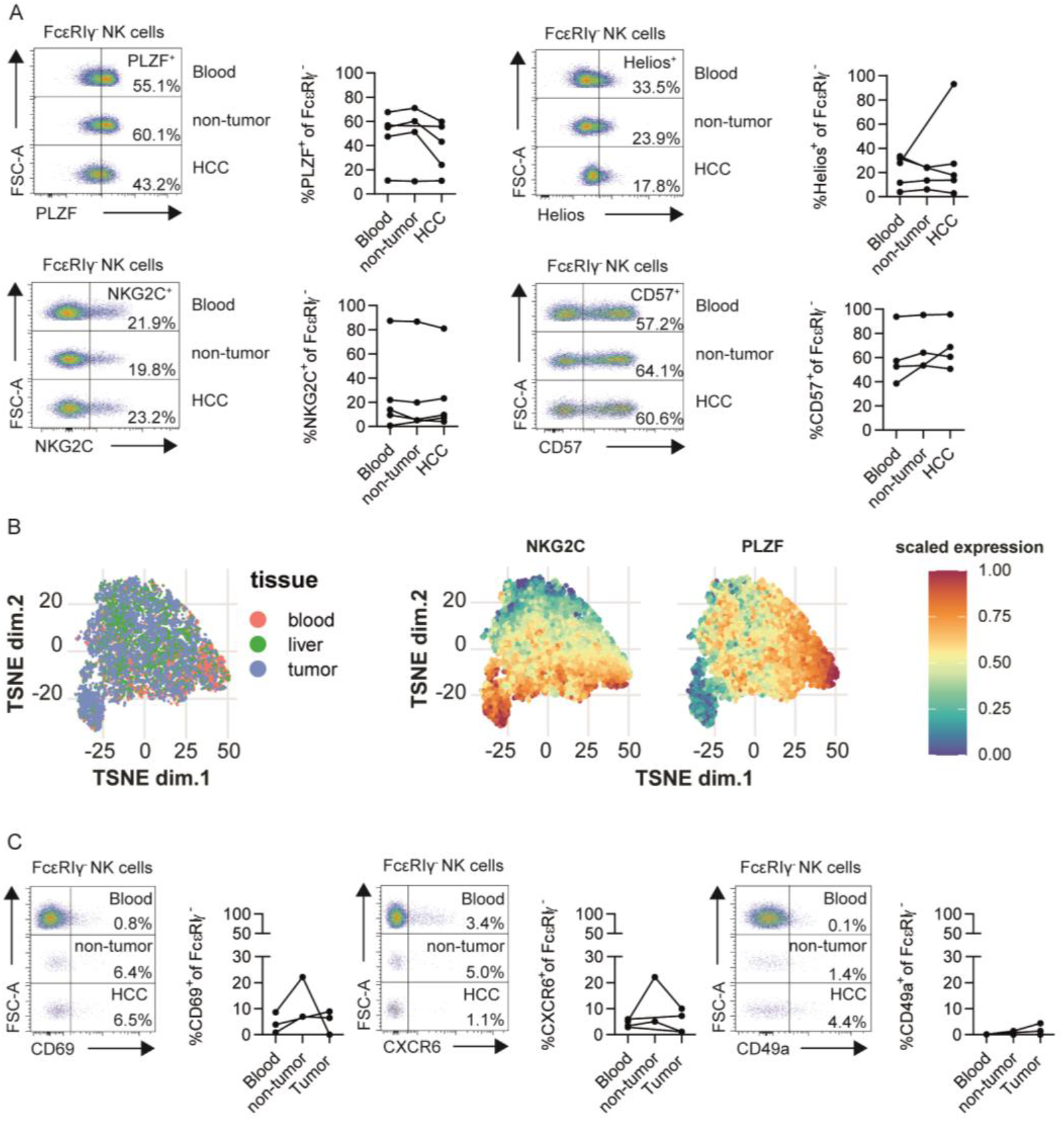
Similar adaptive signature in NK cells isolated from tumor tissue and peripheral blood in HCC patients. PLZF, Helios, NKG2C and CD57 expression of FcεRIγ^-^CD56^dim^ NK cells in blood, adjacent non-tumoral liver and HCC tissue of HCMV^+^ HCC patients (A). t-SNE representation analysis of concatenated flow cytometry data obtained from FcεRIγ^-^CD56^dim^ NK cells in blood, adjacent non-tumoral liver and HCC tissue of HCMV^+^ HCC patients. Expression levels of NKG2C and PLZF are plotted on the t-SNE plots (B). CD69, CXCR6 and CD49a expression of FcεRIγ^-^CD56^dim^ NK cells in blood, non-tumor and HCC tissue of HCMV^+^ HCC patients (C). Each point represents a single HCMV^+^ patient and the samples from one patient are connected by the line. Bars indicate median with IQR. Statistical significance was tested using Kruskal-Wallis test. HCC: hepatocellular carcinoma.

### Limited anti-tumoral activity of FcεRIγ^-^adaptive NK cells

Next, we analyzed the possible impact of HCMV-driven NK-cell diversification on the anti-tumoral activity in HCC. For this, we performed co-culture experiments of FcεRIγ– based NK-cell populations obtained from the blood of HCMV^+^ HCC patients (>10% FcεRIγ^-^adaptive CD56^dim^ NK cells) with the hepatoma cell lines Huh7 and HepG2. CD107a expression, as surrogate marker for degranulation, and MIP-1ß production after stimulation with Huh7 (Fig. 5A, SI Fig.5A and SI Fig.6A) or HepG2 (Fig. 5B, SI Fig. 5B and SI Fig. 6B) cells were significantly reduced in FcεRIγ^-^adaptive NK cells compared to conventional FcεRIγ^+^ NK cells. This observation indicates reduced anti-tumoral activity against liver cancer cells. We also observed the same effect, after stimulation with the leukemia cell line K562 (Fig. 5C, SI Fig. 5C, SI Fig. 6C, SI Fig. 8A), suggesting that adaptive NK cells only have a poor anti-tumoral activity. Next, to determine whether adaptive NK cells could be activated by the cytokines IL-12 and IL-18, potentially secreted by myeloid cells in the tumor microenvironment, we performed cytokine stimulation of FcεRIγ-based NK-cell populations obtained from HCC patients. In agreement with previous studies [30-35] adaptive FcεRIγ^-^NK cells exhibited reduced cytokine responsiveness mirrored by cytokine secretion (SI Fig. 5D, SI Fig. 6D, SI Fig. 8B) [30, 35] and also by diminished degranulation (Fig. 5D). Next, we analyzed the functionality of FcεRIγ–based NK-cell populations after CD16-mediated crosslinking in HCMV^+^ individuals with more than 10% FcεRIγ^-^adaptive CD56^dim^ NK cells. We assessed degranulation (Fig. 5E), IFNγ (SI Fig. 5E) and MIP-1ß (SI Fig. 6E) production. CD16-mediated functionality was comparable between adaptive FcεRIγ^-^CD56^dim^ NK cells compared to conventional FcεRIγ^+^ CD56^dim^ NK cells in HCMV^+^ HCC patients. To analyze the immunoregulatory potential of adaptive FcεRIγ^-^CD56^dim^ NK cells compared to conventional FcεRIγ^+^ CD56^dim^ NK cells towards CD8^+^ T cells, that represent important anti-tumoral effector cells in HCC [2, 5], we co-cultivated FcεRIγ– based NK-cell populations from HCC patients with autologous, activated CD8^+^ T cells. Degranulation (Fig. 5F by FcεRIγ^-^CD56^dim^ NK cells was significantly reduced compared to FcεRIγ^+^ CD56^dim^ NK cells with a similar trend with respect to cytokine production (SI Fig. 5F and SI Fig. 6F). This observation suggests a reduced immunoregulatory activity of adaptive NK cells towards CD8^+^ T cells. Thus, while this may potentially lead to an indirect support of the CD8^+^ T cell-based anti-tumoral response in HCC the direct anti-tumoral activity mediated by adaptive NK cells is limited.

**Fig. 5:**
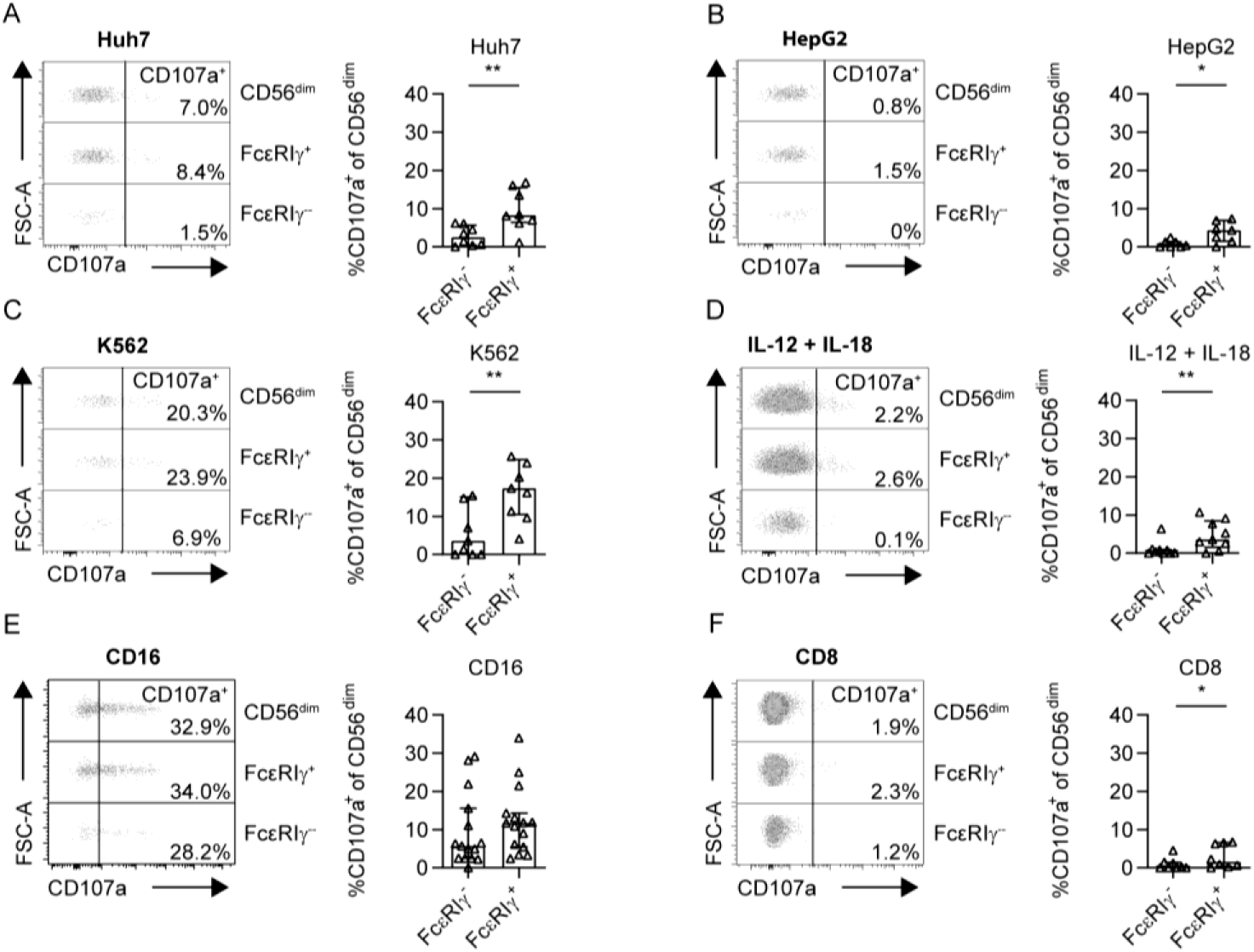
FcεRIγ^-^ adaptive NK cells show reduced antitumor activity compared to conventional NK cells in HCC patients. CD107a expression of CD56^dim^ NK cells following stimulation with Huh7 (A), HepG2 (B) or K562 (C) cell lines for 5h, cytokine stimulation with IL-12 and IL-18 overnight (D) CD16 crosslink (E) or stimulation with activated autologous CD8^+^ T cells for 5h (F) in HCMV^+^ HCC patients. Each dot represents an HCMV^+^ individual. Bars indicate median with IQR. Patients with less than 10% adaptive FcεRIγ^-^CD56^dim^ NK cells were excluded. The following statistical analyses were performed: two-tailed Wilcoxon-Test.

**Fig. 6:**
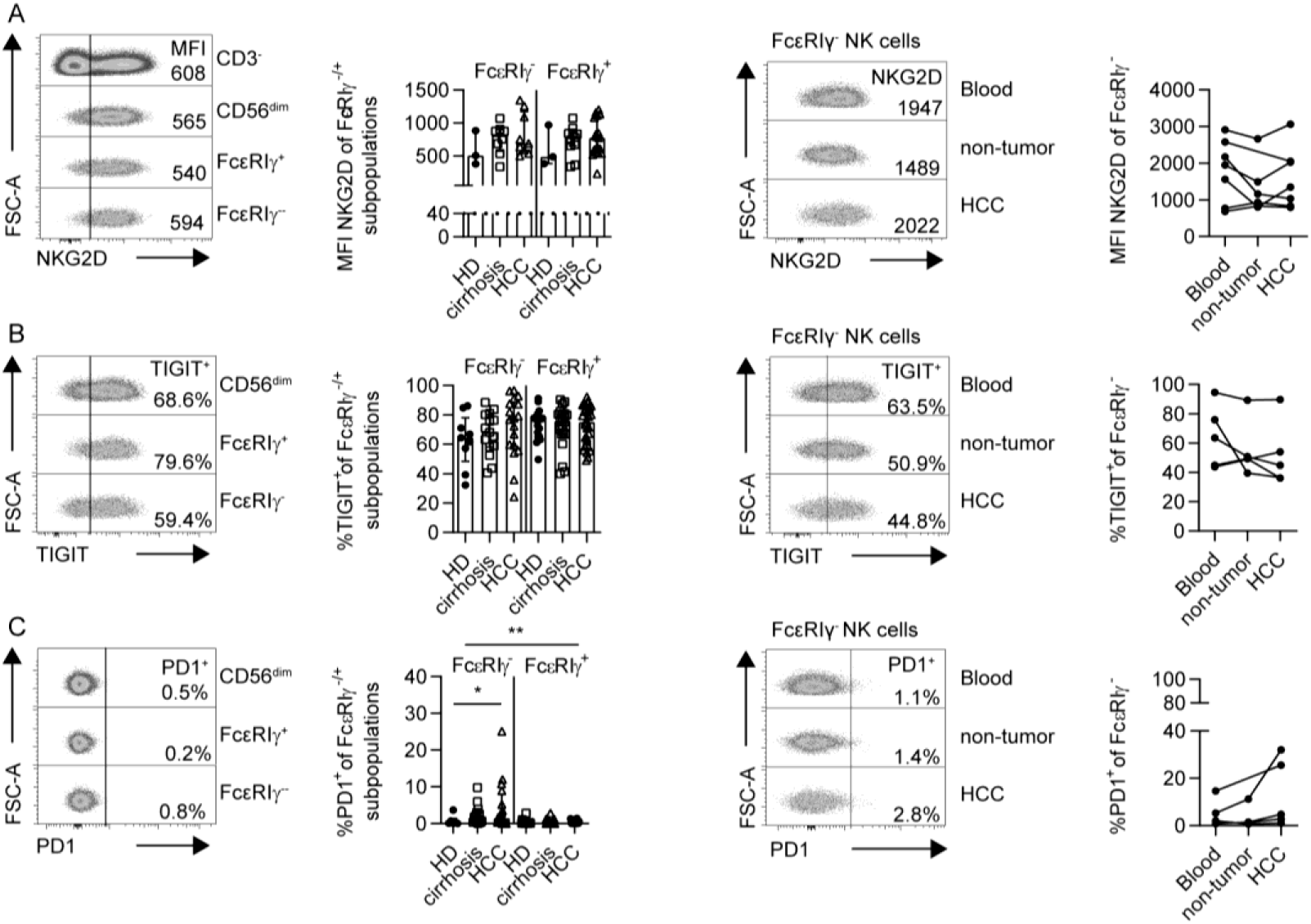
Conserved phenotypic profile of NK cells in HCC patients. MFI of NKG2D (A), expression of TIGIT (B) and PD1 after deduction of FMO (C) of FcεRIγ-subpopulations in HCC patients, HD and patients with liver cirrhosis (left) and in HCC patients’
s blood compared to non-tumor and HCC lesion (right). Each dot represents an HCMV^+^ individual with more than 10% adaptive FcεRIγ^-^CD56^dim^ NK cells and the samples from one patient are connected by the line. Bars indicate median with IQR. Statistical significance was tested by using two-way ANOVA (left) and Kruskal-Wallis test (right). HCC: patients with hepatocellular carcinoma, HD: healthy donors, cirrhosis: patients with liver cirrhosis.

### Adaptive NK cells are not impaired in HCC

Next, we addressed the question whether there is an HCC-associated impairment of adaptive NK cells. For this, we first analyzed the surface expression pattern of the activating receptor NKG2D on FcεRIγ–based CD56^dim^ NK-cell populations that has been shown to mediate altered-self signals from tumors [10, 11]. We did not detect differences in the NKG2D expression neither on adaptive FcεRIγ^-^nor on FcεRIγ^+^ NK cells from HCC patients compared to patients suffering from cirrhosis or HD (Fig. 6A). Next, we analyzed the expression of the so-called checkpoint inhibitory receptors TIGIT and PD1 on FcεRIγ-based CD56^dim^ NK-cell populations. While TIGIT expression on FcεRIγ^-^CD56^dim^ NK cells was comparable to FcεRIγ^+^ CD56^dim^ NK cells (Fig. 6B), a higher fraction of adaptive FcεRIγ^-^CD56^dim^ NK cells expressed PD1 (Fig. 6C). Of note, overall, only very few CD56^dim^ NK cells carried PD1 on their cell surface (Fig. 6C). Nevertheless, PD1 expression was significantly increased on FcεRIγ^-^adaptive CD56^dim^ NK cells from HCMV^+^ HCC patients compared to HCMV^+^ HD. Since this increased expression of PD1 on FcεRIγ^-^CD56^dim^ NK cells from HCMV^+^ HCC patients may be a phenotypic mark of functional impairment, we next tested the functional capacities of FcεRIγ-based CD56^dim^ NK-cell subpopulations from HCMV^+^ HCC patients and from HCMV^+^ cirrhosis patients and HCMV^+^ HD. We did not observe differences in the degranulation (Fig. 7A) and cytokine secretion (SI Fig. 7) of adaptive NK cells obtained from HCMV^+^ HCC patients compared to HCMV^+^ control cohorts, neither after stimulation with tumor cells (K562, HepG2 and Huh7) nor with cytokines (IL-12/IL-18). Degranulation (Fig. 7A) and cytokine production (SI Fig. 7E) after CD16 crosslink did also not differ between HCMV^+^ HCC patients and HCMV^+^ control cohorts. In addition, the immunoregulatory activity was not altered in adaptive NK cells obtained from HCMV^+^ HCC patients compared to HCMV^+^ control cohorts as tested by co-cultivation with activated autologous CD8^+^ T cells (SI Fig. 7F). These data demonstrate that the functional profile of FcεRIγ-based NK-cell subpopulations is conserved in HCMV^+^ HCC and thus, adaptive and conventional CD56^dim^ NK cells are not functionally impaired. However, FcεRIγ^-^adaptive NK cells exhibit altered functional capacities compared to FcεRIγ^+^ conventional CD56^dim^ NK cells. While no significant differences in the NKG2A expression were detected between CD56^dim^ NK cell subpopulations, the lower NCR (NKp30, NKp46) expression (Fig. 7B) in FcεRIγ^-^adaptive CD56^dim^ NK cells compared to FcεRIγ^+^ conventional CD56^dim^ NK cells could possibly explain the differences between the anti-tumoral capacities of CD56^dim^ NK-cell subpopulations.

**Fig. 7:**
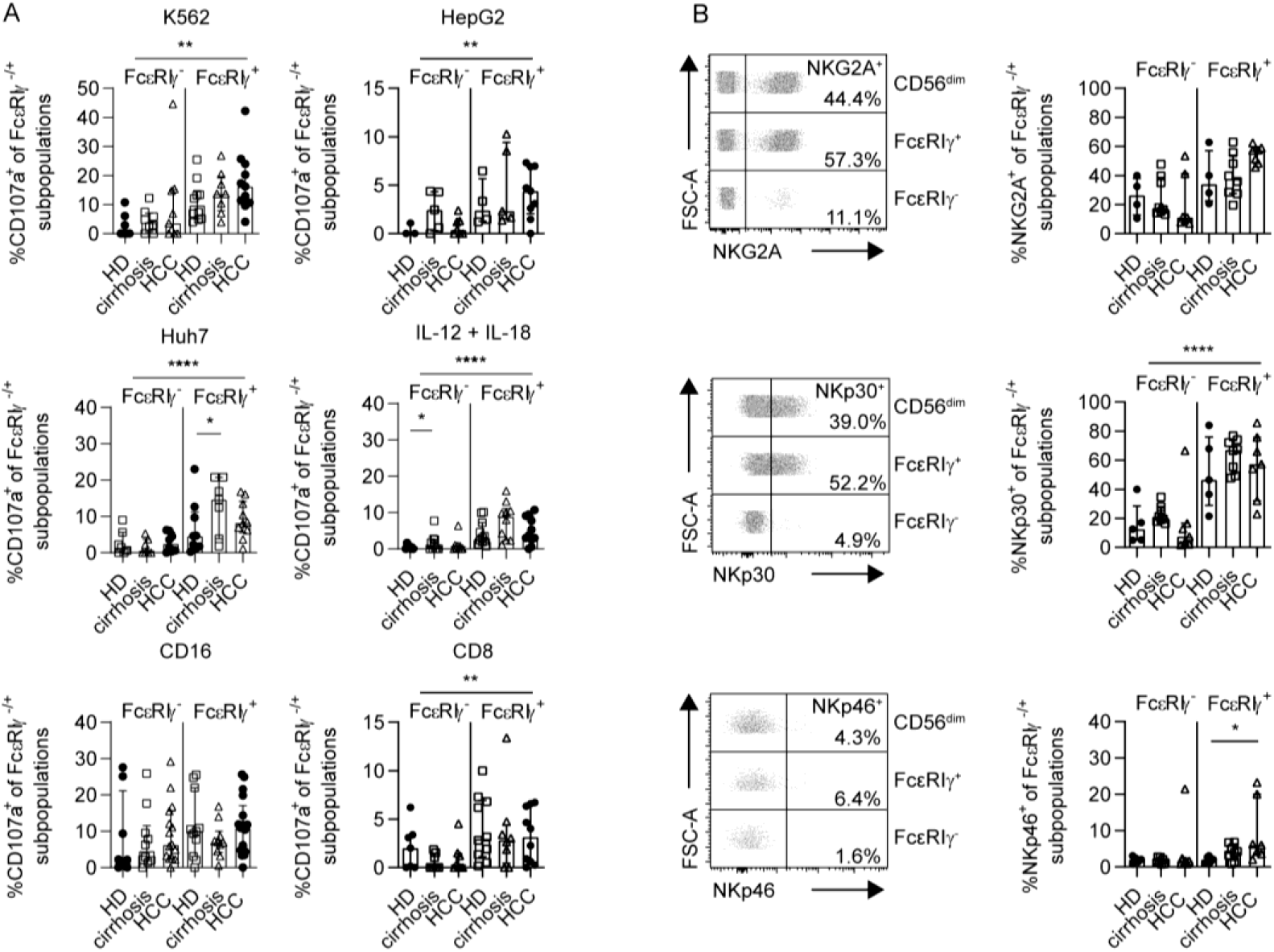
Conserved functionality and receptor repertoire of CD56^dim^ NK cell subpopulations. CD107a expression of FcεRIγ-subpopulations in HCC patients, HD and patients with liver cirrhosis after stimulation with K562, HepG2 and Huh7 cell lines, cytokines, CD16 crosslink or autologous CD8^+^ T cells (A). NKG2A, NKp30 and NKp46 expression of FcεRIγ-subpopulations in HCC patients, HD and patients with liver cirrhosis (B). Each dot represents an HCMV^+^ individual with more than 10% adaptive FcεRIγ^-^CD56^dim^ NK cells. Bars indicate median with IQR. Statistical significance was tested by using two-way ANOVA. HCC: patients with hepatocellular carcinoma, HD: healthy donors, cirrhosis: patients with liver cirrhosis.

**Fig. 8:**
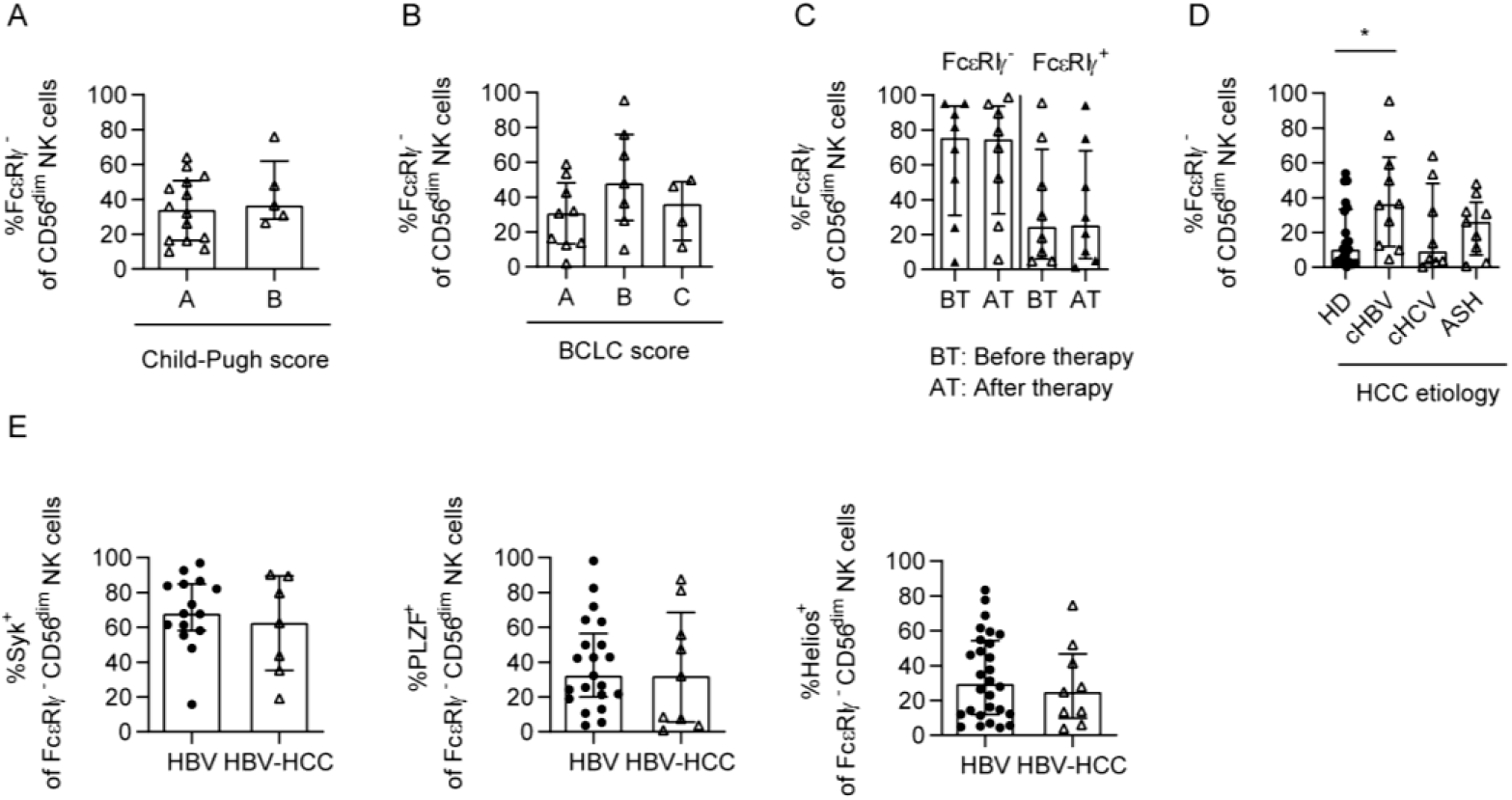
Expansion of FcεRIγ^-^ adaptive NK cells in HBV-associated HCC patients. Frequency of FcεRIγ^-^adaptive NK cells in HCC patients depending on Child scores (A), BCLC scores (B) HCC therapy (C), and underlying etiology of HCC (D). Syk, PLZF and Helios expression on FcεRIγ^-^adaptive CD56^dim^ NK cells in HCMV^+^ HBV infected patients and HCMV^+^ HBV-associated HCC patients (E). Each dot represents an HCMV^+^ individual with more than 10% adaptive FcεRIγ^-^CD56^dim^ NK cells. Bars indicate median with IQR. The following statistical analyses were performed: unpaired, two-tailed Mann-Whitney test (A and E), Kruskal-Wallis test (B and D), paired two-tailed Wilcoxon-Test (C). ASH: alcohol induced steatohepatitis, AT: after therapy, BT: before therapy, HBV: hepatitis B virus, HBV-HCC: HBV-associated HCC patients, HCC: patients with hepatocellular carcinoma, HCMV: human cytomegalovirus, HCV: hepatitis C virus, HD: healthy donors.

### Adaptive NK cells are expanded in HCMV^+^ HBV-associated HCC patients

Based on the altered functional profile of adaptive NK cells, especially the reduced anti-tumoral activity that is conserved in HCC, we wondered whether the frequencies of FcεRIγ^-^CD56^dim^ NK cells are associated with clinical parameters. We did not detect different frequencies of FcεRIγ^-^CD56^dim^ NK cells dependent on the Child-Pugh score, reflecting the severity of cirrhosis and liver function (Fig. 8A), and the BCLC score (Fig. 8B) used to stage HCC. Similarly, no significant correlation of FcεRIγ^-^CD56^dim^ NK-cell frequencies with the tumor marker AFP could be observed (SI Fig. 9). Next, we analyzed whether ablative therapy had an impact on the distribution of FcεRIγ-based NK-cell subpopulations. As shown in Fig. 8C, we did not observe changes in the frequencies of FcεRIγ^+^ or FcεRIγ^-^CD56^dim^ NK cells induced by ablative therapy. Previously, our group showed that higher frequencies of adaptive NK cells are detectable in patients suffering from chronic HBV infection compared to HD [35]. Thus, we next assessed the impact of the underlying HCC etiology, including HBV infection, on the presence of adaptive NK cells in HCMV^+^ HCC patients. Indeed, adaptive NK-cell frequencies were significantly increased in HBV-associated HCMV^+^ HCC patients compared to HCMV^+^ HD but not in HCMV^+^ HCV-or ASH-associated HCC (Fig. 8D). Thus, HBV infection also alters the NK-cell repertoire in HCMV^+^ HCC patients. Of note, Syk, PLZF and Helios (Fig. 8E) expression, cytokine stimulation (SI Fig. 10A) and ADCC (SI Fig. 10B) are similar of NK cells from patients with chronic HBV infection and with HBV-associated HCC. Hence, the etiology is a relevant determinant of the NK-cell repertoire in HCC with implications for the anti-tumoral NK-cell response.

## Discussion

NK cells are potent effector cells in anti-tumoral immunity. In this study, we assessed the impact of CD56^dim^ NK-cell diversification on the NK-cell response in HCC. For this, we performed *ex vivo* analyses of the blood and tumor-derived NK-cell repertoire in HCC patients compared to HD and patients with liver cirrhosis as a state of pre-malignancy.

First, we observed that adaptive CD56^dim^ NK cells were also present in patients suffering from HCC or pre-malignant cirrhosis. The presence of adaptive CD56^dim^ NK cells was clearly linked to HCMV seropositivity as previously reported in other patient cohorts including cohorts affected by viral infections or leukemia [30-33, 35, 36, 38-40]. Since HCMV seropositivity is frequent [41, 42], as also reflected by our cohorts of patients suffering from HCC and cirrhosis, CMV-associated emergence of adaptive CD56^dim^ NK cells is an important parameter of NK cell-mediated cancer immunosurveillance with potential clinical impact.

Second, besides this link to HCMV, we detected increased frequencies of adaptive CD56^dim^ NK cells in HBV-associated HCC compared to the other analyzed etiologies such as HCV and ASH. Furthermore, no other tested clinical parameters including liver function or tumor stage correlated with the frequencies of adaptive CD56^dim^ NK cells arguing against a general HCMV reactivation in HCC lesions accompanied by an expansion of adaptive CD56^dim^ NK cells. The underlying mechanisms of the pronounced expansion of adaptive CD56^dim^ NK cells seem to be rather HBV-related. This is in line with a recent report from our group showing increased frequencies of adaptive CD56^dim^ NK cells in patients chronically infected with HBV compared to HCV and HD [35]. However, further studies are required to uncover these HBV-related mechanisms. Possible mechanisms may include the presence of anti-HCMV/anti-HBV antibodies, selection for specific HCMV strains, cytokines and also genetic factors [35]. Third, adaptive CD56^dim^ NK cells were not only present in the blood but also detectable in liver tissue, non-HCC as well as HCC-derived. Thus, this subset is not only involved in NK cell-mediated cancer immunosurveillance in blood-born malignancies [34, 40] or in metastasis but also within solid tumor tissue like HCC. Interestingly, we observed similar frequencies of adaptive NK-cell subsets within the circulation and in tumor and adjacent non-tumor liver tissue. This indicates that adaptive CD56^dim^ NK cells rather belong to infiltrating lymphocytes instead of being tissue-resident. This is in line with our findings that adaptive CD56^dim^ NK cells displayed a similar phenotype in blood and tumor and that non-HCC- and HCC-derived adaptive NK-cell subsets exhibited only a low expression of tissue-resident markers [19, 27, 28, 43], like CXCR6, CD69 and CD49a. Hence, adaptive CD56^dim^ NK cells found within HCC tissue most probably represent circulating/tumor-infiltrating lymphocytes that can be monitored and targeted via the blood. This observation may have important relevance for translational applications, such as for immunotherapeutic interventions. However, additional studies are required to further analyze the circulating/infiltrating nature of adaptive CD56^dim^ NK cells within the liver, e.g. by the analysis of the KIR repertoire to exclude that adaptive NK cells are locally expanded.

Forth, compared to conventional FcεRIγ^+^ CD56^dim^ NK-cell subset, adaptive FcεRIγ^-^CD56^dim^ NK cells displayed reduced direct anti-tumoral activity. This limited anti-tumoral activity was not only evident against a leukemia cell line, as reported previously [34], but also against hepatoma cells, indicating a rather inherently reduced response triggered by low MHCI expression. This limited response of adaptive CD56^dim^ NK cells towards MHCI^low^ target cells may be detrimental in cancer immunosurveillance reflecting a possible successful cancer evasion strategy. However, it also potentially represents a mechanism of tissue protection, especially in the liver since hepatocytes express low levels of MHCI [44]. A general underlying mechanism is further supported by the observation that adaptive CD56^dim^ NK cells from HCC patients and HD exhibited a similarly limited anti-tumoral activity. Thus, it is not specifically induced by the HCC microenvironment but it is rather an intrinsic and stable characteristic of adaptive CD56^dim^ NK cells. This also fits to our data regarding an unaltered expression of NKG2D, TIGIT and NKG2A on adaptive CD56^dim^ NK cells obtained from HCC patients compared to HD. These molecules have been reported to be important in anti-tumoral responses mediated by NK cells in which NKG2D acts as activating receptor by binding to altered stress ligands on tumor cells [10, 11] and TIGIT and NKG2A are checkpoint inhibitors [45, 46].

Expression of PD1, another checkpoint inhibitor molecule, was also described on CD56^dim^ NK cells of cancer patients and healthy controls [47, 48]. In particular, Liu *et al*. reported elevated PD1 expression on NK cells obtained from HCC patients [48]. Furthermore, in another study analyzing HCMV seropositive individuals, PD1^+^ NK cells exhibited a higher expression of CD57 [47], a maturation marker that is also highly expressed on adaptive CD56^dim^ NK cells [30]. In line with these reports, we detected an increased expression of the checkpoint inhibitor PD1 on adaptive compared to conventional CD56^dim^ NK cells that was even significantly higher in HCC patients compared to HD. However, the overall frequency of PD1 expressing CD56^dim^ NK cells was extremely low, in the blood, adjacent non-tumor and tumor liver tissue of HCC patients. Hence, we assume a minor role for this checkpoint inhibitor molecule in adaptive CD56^dim^ NK cell-mediated immunosurveillance in HCC [49]. Of note, tumor-infiltrating NK cells in mice and colon cancer patients also showed low expression rates of PD1 [45] questioning a direct role of PD1 as checkpoint inhibitor of NK cells in general.

In addition to the limited direct anti-tumoral activity, adaptive CD56^dim^ NK cells obtained from HCC patients displayed reduced cytokine responsiveness and decreased immunoregulatory function towards activated CD8^+^ T cells. The decreased immunoregulatory function of adaptive CD56^dim^ NK cells may result from the low NCR expression as it has also been previously described in the LCMV mouse system [50]. This functional profile of adaptive CD56^dim^ NK cells was similar in HCC and control groups. These previously described functional features of adaptive NK-cell subsets [30] therefore seem to be conserved in the HCC microenvironment. In the context of cancer immunosurveillance these functional features may favor tumor-specific immunity by T cells on the expense of a broad, less specific anti-tumoral defense induced by altered-self, missing-self and bystander/cytokine-mediated mechanisms. In sum, HCMV-associated CD56^dim^ NK-cell diversification into adaptive and conventional subsets affects cancer immunosurveillance in HCC due to generally imprinted features of the respective subsets. This is in contrast to the HCC-associated dysfunction observed within the liver-resident CD56^bright^ NK-cell population that is at least in part directly induced by HCC in a contact-dependent manner [20]. Based on the observation that adaptive CD56^dim^ NK cells are more frequent in HBV-associated HCC, the limited anti-tumoral activity of this subset becomes especially important in this context and therefore suggests that HCC etiology is linked to differences in cancer immunosurveillance, a fact that has to be considered in the design and application of immunotherapies in HCC.

## Methods

### Study cohort

57 HCC patients (SI table 1), 36 healthy donors (SI table 2) and 33 patients with liver cirrhosis (SI table 3) were recruited at the Department of Medicine II of the University Hospital Freiburg, Germany. Written informed consent was obtained in all cases and the study was conducted according to the Declaration of Helsinki (1975), federal guidelines and local ethics committee regulations (Albert-Ludwigs-University, Freiburg, Germany, approvals 474/14 and 152/17). HCC and adjacent non-tumor liver tissue samples were obtained from the University Hospital Freiburg and non-tumoral liver samples from the Karolinska University Hospital, Stockholm.

### PBMC isolation

Peripheral blood mononuclear cells (PBMCs) were isolated from EDTA anti-coagulated blood by density-gradient centrifugation. Frozen PBMCs were thawed in complete RPMI culture medium (RPMI 1640 supplemented with 10% fetal bovine serum, 1% penicillin/streptomycin and 1.5% 1M HEPES (all Thermo Fisher, USA)).

### Isolation of lymphocytes from tissue samples

Lymphocytes were isolated from tissue samples by density-gradient centrifugation after mechanical dissociation in complete RPMI culture medium (RPMI 1640 supplemented with 10% fetal bovine serum, 1% penicillin/streptomycin and 1.5% 1M HEPES (all Thermo Fisher, USA)).

### HCMV status

HCMV serostatus was determined via HCMV-IgG Chemiluminescence Immunoassay (DiaSorin LIAISON) by the Department of Virology, University of Freiburg or as previously described by Marquardt *et al*. [43]. Briefly, 10^6^ cells were stimulated with 1µg/mL CMVpp65 overlapping peptides (JPT Peptide Technologies, Germany) in the presence of Brefeldin A (GolgiPlug (final concentration 5 µL/mL, BD Biosciences, USA) and Monensin (GolgiStop (final concentration 3.25 µl/ml, BD Biosciences, USA). After 16 h, cytokine production was assessed by flow cytometry [43]. HCMV seronegative (HCMV^-^) individuals are only included in Fig. 1.

### Assessment of NK-cell function

Degranulation and cytokine production of NK cell subsets were determined after cytokine stimulation o/n with IL-12 (10ng/ml, Sigma Aldrich, USA) and IL-18 (5ng/ml, MBL, USA), after CD16-crosslinking or after co-culture with the cell lines K562, Huh7 and HepG2 and autologous CD8^+^ T cells (pre-activated with ImmunoCult™ Human CD3/CD28 T Cell Activator, Stemcell Technologies, Canada according to the manufacturer’s instruction). Detailed information is provided in the supplementary materials and methods.

### Multiparametric flow cytometry

Detailed list of antibodies and reagents used for flow cytometry analysis is included in the supplemental information.

### Statistics

Statistical analysis was performed with GraphPad Prism 6 (GraphPad Software, USA). Statistical tests used are indicated in the figure legends. Levels of significance are indicated as follows: *: *P* <0.05; **: *P* <0.01; ***: *P* <0.001; ****: *P* <0.0001.

## Abbreviations

HCC: hepatocellular carcinoma
NK cell: natural killer cell
HCMV: human cytomegalovirus
HD: healthy donors
HBV: hepatitis B virus
HCV: hepatitis C virus
ADCC: antibody-dependent cell-mediated cytotoxicity
PD1: programmed death receptor
1AFP: α-fetoprotein
ASH: alcohol induced steatohepatis
cirrhosis: patients with liver cirrhosis
IQR: interquartile range
MFI: mean fluorescence intensity
FMO: fluorescence minus one

## Acknowledgements

We thank all patients and donors for participating in the current study. We thank Daniela Huzly and her team for testing HCMV serology.

