## Supplemental Information for "Adaptive subsets limit the anti-tumoral NK-cell activity in Hepatocellular Carcinoma"

### Supplemental materials and methods

#### *Assessment of NK-cell function*

For cytokine stimulation, $10^6$ PBMCs were incubated overnight (o/n) at 37°C in IMDM culture medium (IMDM (Gibco-ThermoFisher), 10% fetal bovine serum, 1% penicillin-streptomycin and 50µM β-mercaptoethanol (Sigma-Aldrich) in the presence of IL-12 and IL-18 (IL-12 10ng/mL, Sigma-Aldrich; IL-18 5ng/mL, MBL) or without cytokines as negative control. Anti-CD107a mAb and anti-CD56 mAb were added directly. 0.96nmol/ml Brefeldin A (BD Biosciences) and 0.25nmol/ml Monensin (BD Biosciences) were added for the last four hours of incubation.

For co-culture assays with the cell lines K562, HuH7 and HepG2 cells NK cells were isolated from PBMCs using MACS cell separation technology (NK cell isolation kit, Miltenyi Biotech) according to the manufacturer's instructions. Isolated NK cells and target cells or NK cells alone as a negative control were incubated in IMDM culture medium in the presence of anti-CD107a mAb and anti-CD56 mAb for five hours at 37°C. 0.96nmol/ml Brefeldin A (BD Biosciences) and 0.25nmol/ml Monensin (BD Biosciences) were added for the last four hours of stimulation. The effector to target ratio was 1:5 for stimulation with HuH7 and HepG2h cell lines and 1:10 for stimulation with K562 cell line.

For CD16-crosslinking, high-binding 96-well flat-bottom plates (Sigma-Aldrich) were coated overnight (o/n) at 4°C with anti-CD16 pure IgG (10µg/ml in PBS, BD Bioscience) or PBS only, as negative control. After extensive washing, $10^6$ PBMCs in IMDM culture medium in the presence of anti-CD107a mAb and anti-CD56 mAb were added to the wells and incubated for five hours at 37°C and 5% $CO_2$. 0.96nmol/ml Brefeldin A (BD Biosciences) and 0.25nmol/ml Monensin (BD Biosciences) were added for the last four hours of incubation (37°C and 5% $CO_2$).

For stimulation with autologous activated $CD8^+$ T cells, $CD8^+$ T cells were isolated from PBMCs using MACS cell separation technology ($CD8^+$ T cell isolation kit, Miltenyi Biotech) according to the manufacturer's instructions. Isolated $CD8^+$ T cells were stimulated for 60h with ImmunoCult™ Human CD3/CD28 T Cell Activator, (Stemcell Technologies) in complete RPMI culture medium. Activated $CD8^+$ T cells and freshly isolated autologous NK cells were co-incubated at a T-cell/NK-cell ratio of

1:10 for 5h in the presence of anti-CD107a mAb and anti-CD56 mAb. 0.96 nmol/ml Brefeldin A (BD Biosciences) and 0.25 nmol/ml Monensin (BD Biosciences) were added for the last four hours of stimulation. NK cells were isolated from PBMCs using MACS cell separation technology (NK cell isolation kit, Miltenyi Biotech) according to the manufacturer's instructions.

Subsequently, surface and intracellular stainings were performed.

***Multiparametric flow cytometry***

For flow cytometry the following antibodies were used: anti-CD3 (SK7), anti-CD4 (SK3), anti-CD8 (RPA-T8), anti-CD16 (3G8), anti-CD49a (SR84), anti-CD56 (B159), anti-CD57 (NK-1), anti-CD107a (H4A3), anti-CXCR6 (13B1E5), anti-PD1 (EH12.1), anti-PLZF (R17-809), anti-NKp46 (9E2/NKp46), anti-NKp30 (p30-15), anti-CD2 (RPA-2.10) (BD Biosciencesanti-CD3 (SK7 and UCHT1), anti-CD45 (HI30), anti-CD56 (5.1H11), anti-CD57 (QA17A04), anti-CXCR6 (K041E5), anti-IFN-γ (4S.B3), anti-NKG2D (1D11), anti-TIGIT (A15153G) anti-TNF (Mab11), anti-Siglec7 (6-434), anti-CD7 (CD7-6B7), anti-CXCR3 (G025H7) (BioLegend), anti-CD14 (61D3), anti-CD19 (HiB19), anti-CD69 (FN50), anti-Helios (22F6), anti-MIP-1β (FL34Z3L), anti-Syk (4D10.1), anti-TIGIT (MBSA43), anti-CX3CR1 (2A9-1) (eBioscience-Thermo) anti-FcεRIγ (polyclonal) (Millipore), anti-NKG2A (131411) and anti-NKG2C (134591) (R&D Systems).

Fixable Viability Dyes (eFluor780, eBioscience-Thermo Fisher) was used for live/dead discrimination. FoxP3/Transcription Factor Staining Buffer Set (eBioscience-Thermo Fisher) was applied according to the manufacturer's instructions for intranuclear staining. Cells were fixed with paraformaldehyde and analyzed using FACSCanto II or LSRFortessa (BD Biosciences).

***Dimensionality reduction of multiparametric flow cytometry data***

The visualization of multiparametric flow cytometry data was done with R using the Bioconductor (CATALYST package (Crowell H, Zanotelli V, Chevrier S, Robinson M (2020). CATALYST: Cytometry dATa anALYSis Tools. R package version 1.12.2, https://github.com/HelenaLC/CATALYST). The analyses were performed on gated FcεRIγ$^{-}$ CD56$^{dim}$ NK cells and included the markers NKG2C, PLZF, CD57, Helios, CD16, PD-1, TIGIT and NKG2D. Down sampling of cells to the group comprising the lowest cell count or to 3000 cells was performed prior to dimensionality reduction in order to facilitate the visualization of different samples. Marker intensities were transformed by arcsinh (inverse hyperbolic sine) with a cofactor of 150. Dimensionality reduction on the transformed data was achieved by t-distributed stochastic neighbor embedding (t-SNE) using the CATALYST package functions runDR with default parameters.

### Supplement figures

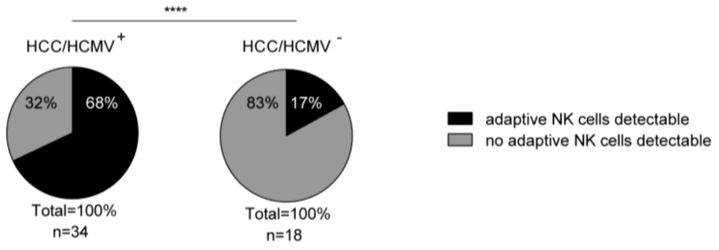

**SI Fig. 1. Increased frequency of FcεRIγ$^-$ NK cells in HCMV$^+$ HCC patients.**

Pie charts depicting presence of with FcεRIγ$^-$ adaptive CD56$^{dim}$ NK cells (black) and absence of with FcεRIγ$^-$ CD56$^{dim}$ adaptive NK cells (grey) in HCMV$^+$ (left) and HCMV$^-$ (right) HCC patients. Patients with adaptive NK cells are defined as >10% FcεRIγ$^-$ adaptive CD56$^{dim}$ NK cells. Statistical analysis was performed with parts of whole analysis (binomial test).

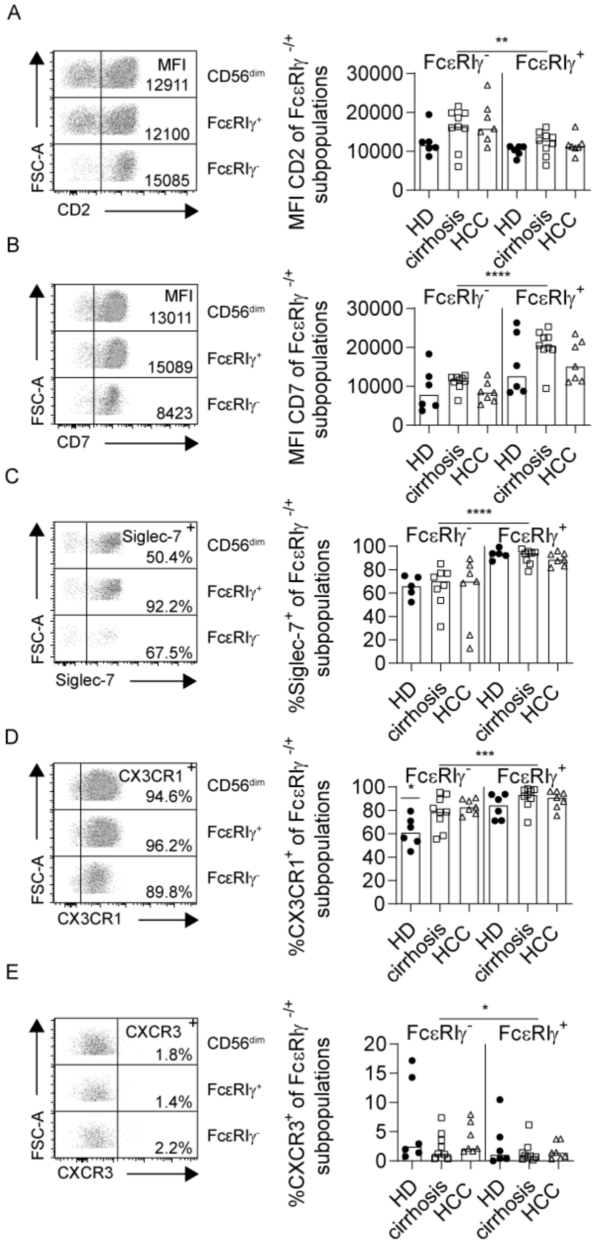

**SI Fig. 2. Adaptive NK-cell profile is comparable between HCC patients and control cohorts.** CD2 (A), CD7 (B), Siglec-7 (C), CX3CR1 (D) and CXCR3 (E) expression on FcεRIγ- subpopulations in HCMV+ HD, HCMV+ patients with liver cirrhosis and HCMV+ HCC patients. Each point represents a single patient with more than 10% adaptive FcεRIγ- of CD56dim NK cells. Bars indicate median. Statistical significance was tested using two-way ANOVA.

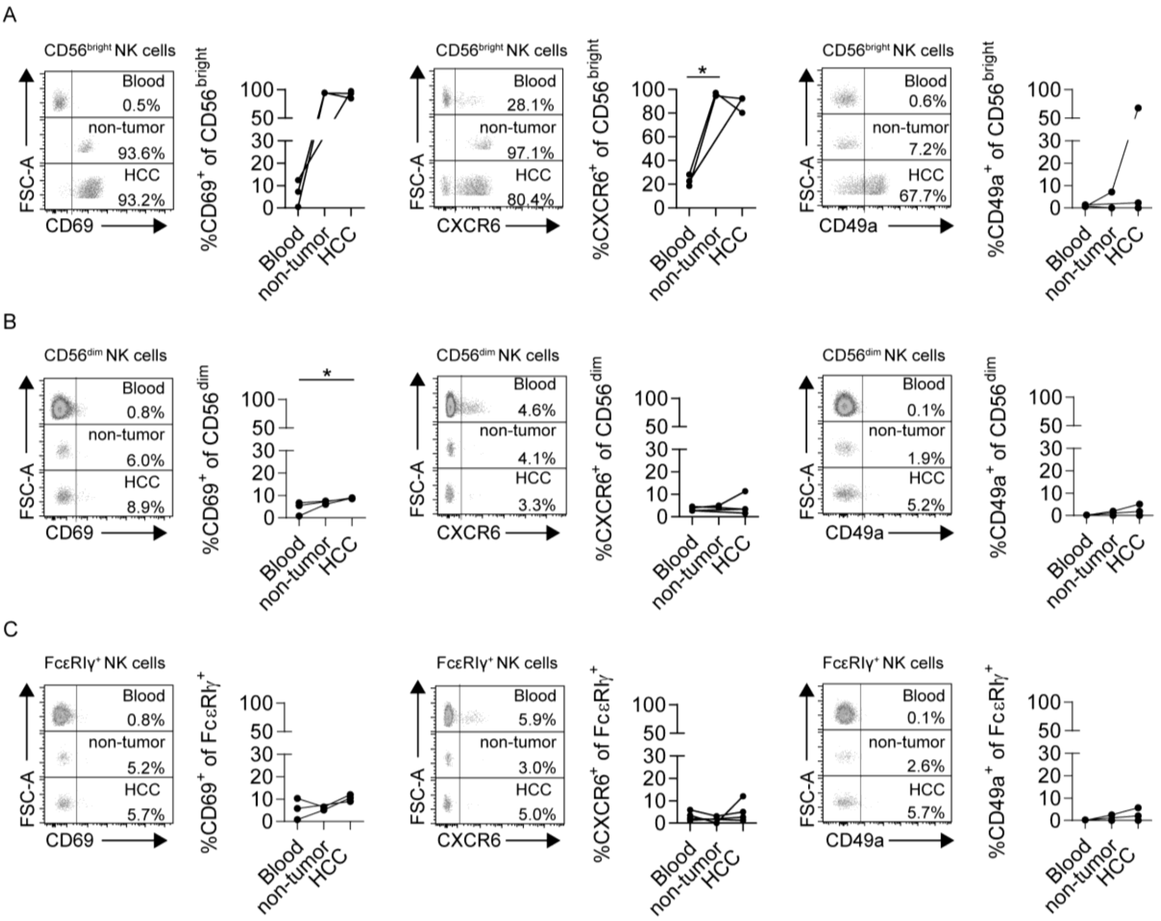

**SI Fig. 3. CD56$^{dim}$ NK cells and subpopulations do not express tissue resident markers in HCC tissue.**

CD69, CXCR6 and CD49a expression on CD56$^{bright}$ (A) CD56$^{dim}$ (B) and FcεRIγ$^{+}$ (C) CD56$^{dim}$ NK cells in blood, non-tumor and HCC tissue of HCC patients. Each point represents a single HCMV$^{+}$ patient and the samples from one patient are connected by the line. Statistical analysis was performed with Kruskal Wallis test.

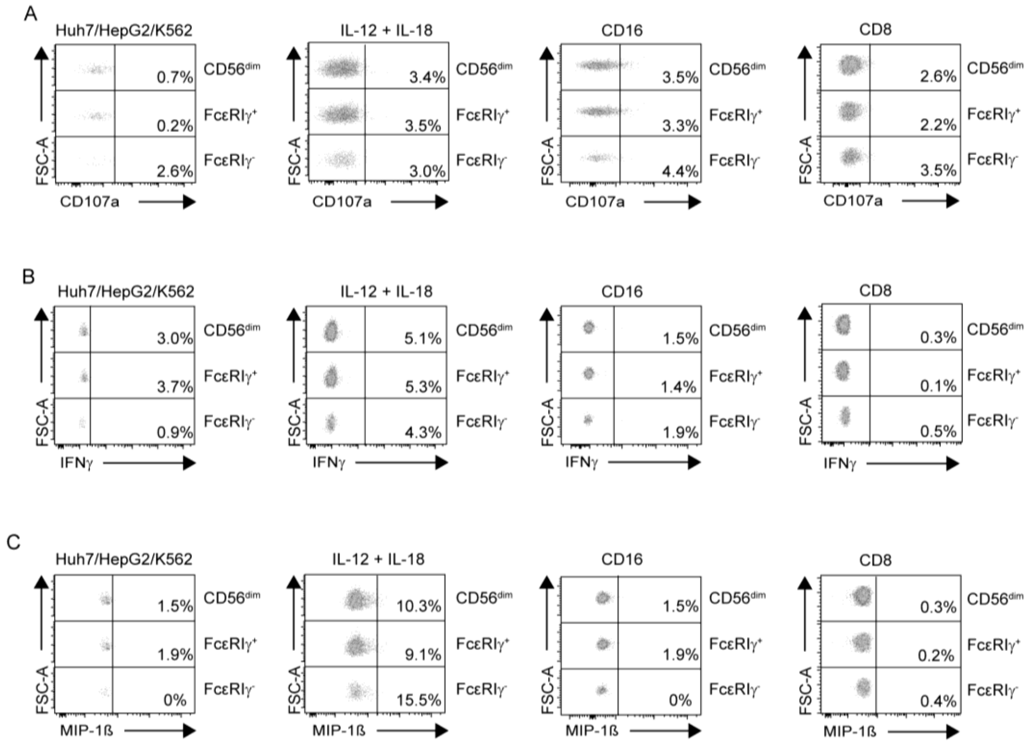

**SI Fig. 4. Negative controls for functional studies.**

CD107a (A), IFNγ (B) and MIP-1ß (C) expression of CD56$^{dim}$ NK cells, FcεRIγ$^{+}$ and FcεRIγ$^{-}$ CD56$^{dim}$ NK cells without stimulation.

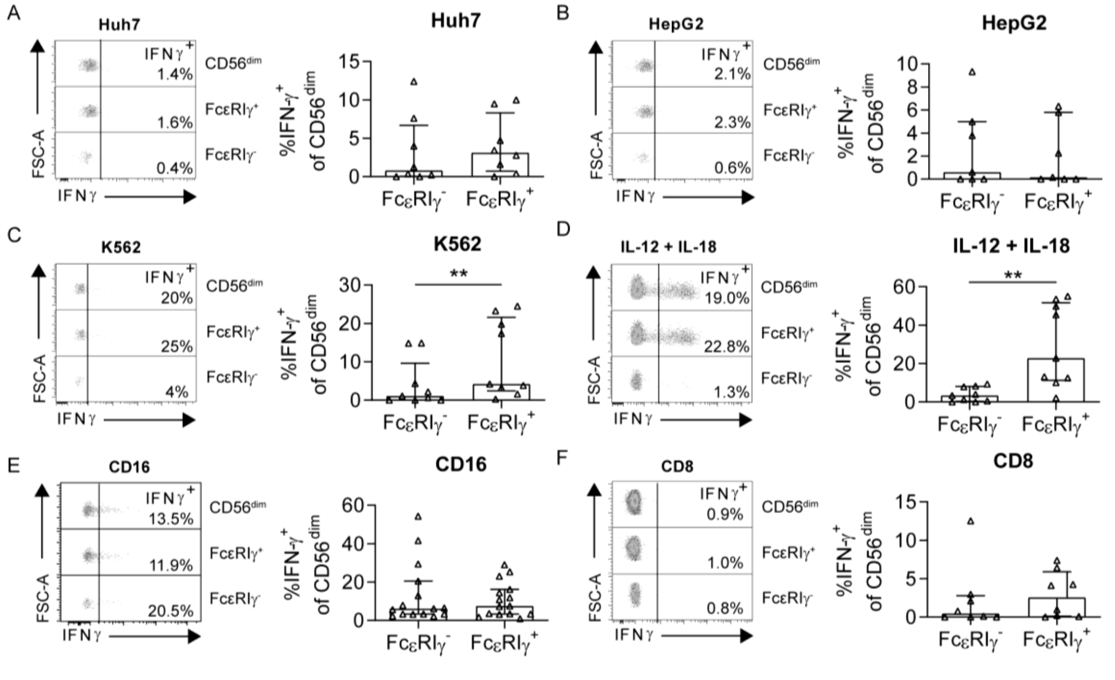

**SI Fig. 5. FcεRIγ$^{-}$ adaptive NK cells show reduced anti-tumoral activity (measured by IFNγ expression) compared to conventional NK cells in HCC patients.**

IFNγ expression of CD56$^{dim}$ NK cells following stimulation with Huh7 (A), HepG2 (B) or K562 (C) cell lines for 5h, cytokine stimulation with IL-12 and IL-18 overnight (D), CD16 crosslink (E) or stimulation with autologous activated CD8$^{+}$ T cells for 5h (F) in HCMV$^{+}$ HCC patients. Each dot represents an individual with more than 10% FcεRIγ⁻ adaptive CD56$^{dim}$ NK cells. Bars indicate median with IQR. Statistical significance was tested using paired two-tailed Wilcoxon test.

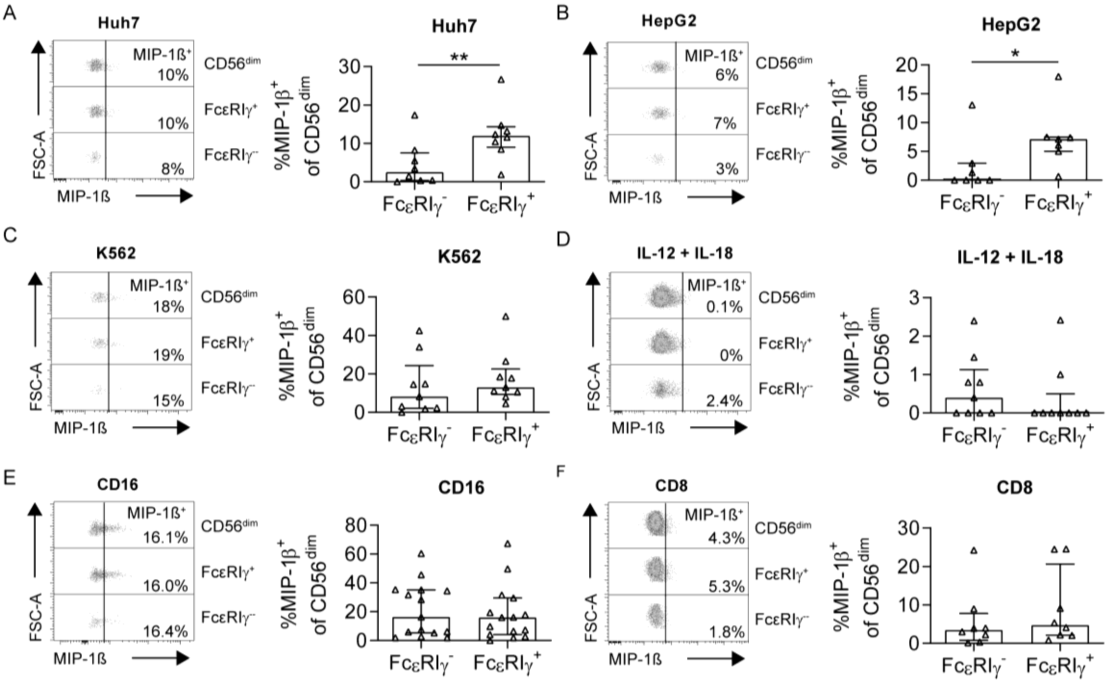

**SI Fig. 6. FcεRIγ⁻adaptive NK cells show reduced anti-tumoral activity (measured by MIP-1ß expression) compared to conventional NK cells in HCC patients.**

MIP-1ß expression of CD56$^{dim}$ NK cells following stimulation with Huh7 (A), HepG2 (B) or K562 (C) cell lines for 5h, cytokine stimulation with IL-12 and IL-18 overnight (D), CD16 crosslink (E) or stimulation with autologous activated CD8$^{+}$ T cells for 5h (F) in HCMV$^{+}$ HCC patients. Each dot represents an individual with more than 10% FcεRIγ⁻ adaptive CD56$^{dim}$ NK cells. Bars indicate median with IQR. Statistical significance was tested using paired two-tailed Wilcoxon test.

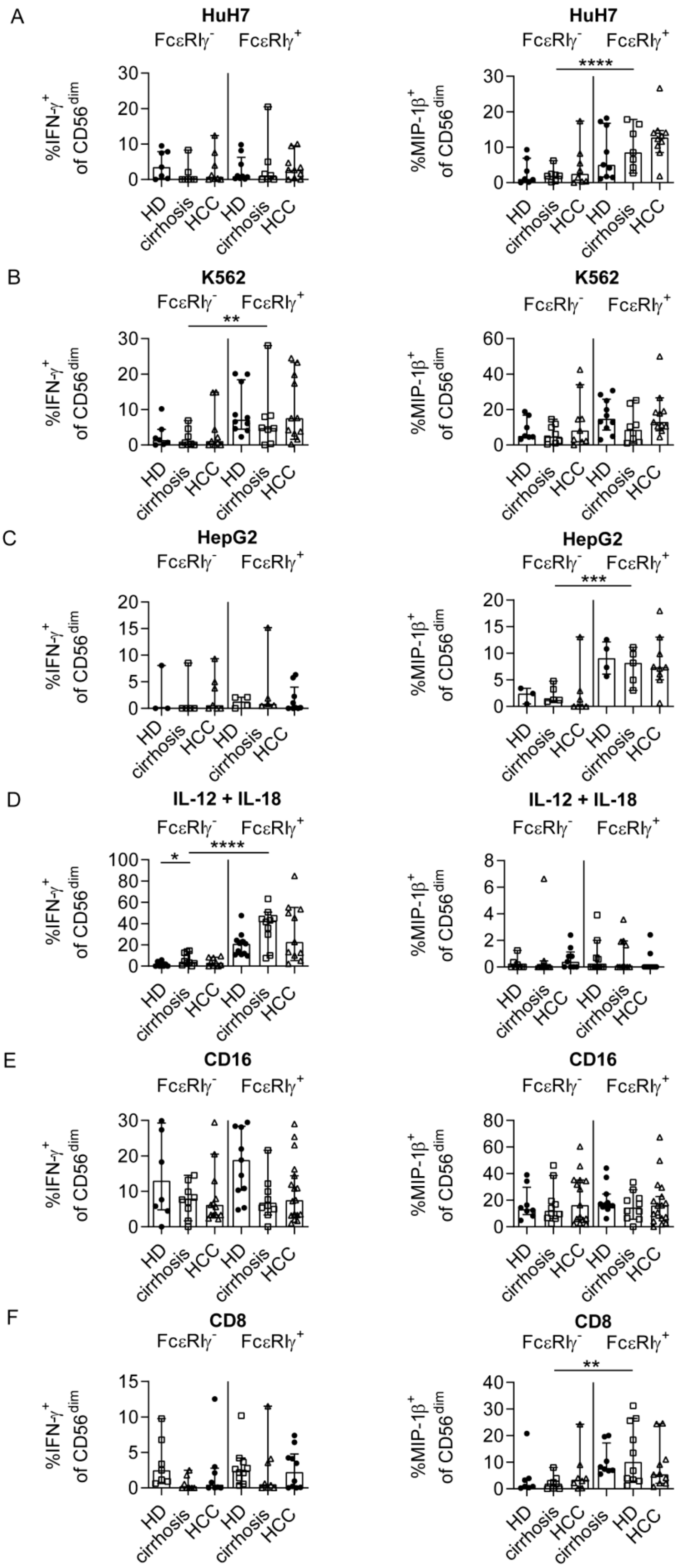
A
HuH7
FcεRIγ⁻ FcεRIγ⁺
%IFN-γ⁺ of CD56dim
%MIP-1β⁺ of CD56dim
****
HD
cirrhosis
HCC
B
K562
**
C
HepG2
***
D
IL-12 + IL-18
*
****
E
CD16
F
CD8
**

**SI Fig. 7. FcεRIγ$^-$ adaptive NK cells show reduced cytokine response compared to conventional NK cells.**

IFNγ and MIP-1ß expression of CD56$^{dim}$ NK cells following stimulation with Huh7 (A), K562 (B) and HepG2 cell lines (C) for 5h, cytokine stimulation with IL-12 and IL-18 overnight (D), CD16 crosslink (E) or stimulation with autologous activated CD8$^+$ T cells for 5h (F) in HCMV$^+$ HCC patients and control cohorts. Each dot represents an individual with more than 10% FcεRIγ$^-$ adaptive CD56$^{dim}$ NK cells. Bars indicate median with IQR. Statistical significance was tested using paired two-way ANOVA. HCMV: human cytomegalovirus, HCC: patients with hepatocellular carcinoma, HD: healthy donors, cirrhosis: patients with liver cirrhosis.

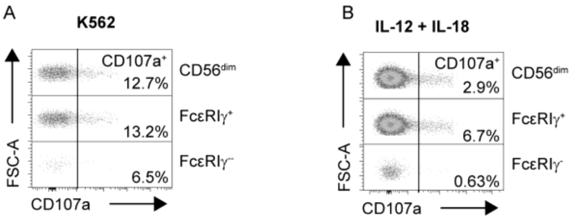

**SI Fig. 8. Reduced anti-tumoral activity of FcεRIγ$^-$ adaptive NK cells compared to conventional NK cells in healthy liver tissue.**

Representative CD107a expression of FcεRIγ$^+$ and FcεRIγ$^-$ CD56$^{dim}$ NK cells from healthy liver tissue after stimulation with K562 cell line for 5 h (A) or cytokine stimulation (IL-12+IL-18) overnight (B) in a donor with more than 10% FcεRIγ$^-$ adaptive CD56$^{dim}$ NK cells.

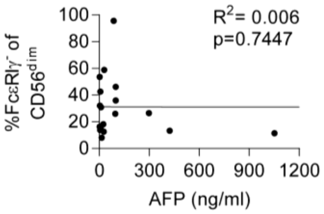

**SI Fig. 9. Frequency of FcεRIγ$^-$ adaptive NK cells do not correlate with AFP values.**

Correlation analysis of AFP value and frequencies of FcεRIγ$^-$ CD56$^{dim}$ NK cells in HCMV$^+$ HCC patients. Statistical analysis was performed with linear regression analysis. Patients with less than 10% FcεRIγ$^-$ adaptive CD56$^{dim}$ NK cells were excluded. HCMV: human cytomegalovirus, HCC: patients with hepatocellular carcinoma.

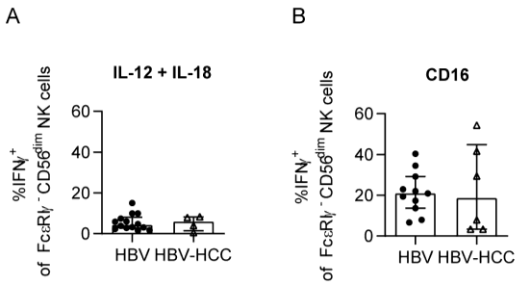

**SI Fig. 10. Conserved functional profile of FcεRIγ$^{-}$ adaptive CD56$^{dim}$ NK cells between HCMV$^{+}$ HBV and HCMV$^{+}$ HBV-associated HCC patients.**

IFNγ expression of CD56$^{dim}$ NK cells following stimulation with cytokine stimulation with IL-12 and IL-18 overnight (A) or CD16 crosslink (B) in HCMV$^{+}$ chronic HBV and HBV-associated HCC patients. Each dot represents an individual with more than 10% FcεRIγ$^{-}$ adaptive CD56$^{dim}$ NK cells. Bars indicate median with IQR. Statistical analysis was performed with Mann-Whitney test. HBV: hepatitis B virus, HCC: patients with hepatocellular carcinoma, HCMV: human cytomegalovirus.

**Supplemental Tables**

**SI Table 1. Study cohort of HCC patients.**

M: male, F: female, AIH: autoimmune hepatitis, ASH: alcohol induced steatohepatis, HBV: hepatitis B virus, HCV: hepatitis C virus, HDV: hepatitis D virus, HFE: hemochromatosis, NASH: non-alcoholic steatohepatis, n.d.: not determined, neg: negative, pos: positive, * NK cells isolated from peripheral blood, non-tumoral liver tissue and HCC lesions; data are depicted in Fig. 3 and 4.

| Age [years] | Sex | Etiology | Child score | BCLC score | AFP [ng/ml] | GPT [U/l] | GOT [U/l] | HCMV sero-status | Treatment prior inclusion | Pheno-typic analy-sis | Funct-ional analy-sis |
|---|---|---|---|---|---|---|---|---|---|---|---|
| 78 | M | ASH | B | B | n.d. | n.d. | n.d. | pos | no | yes | yes |
| 45 | M | HBV | A | C | n.d. | 296 | 235 | pos | no | yes | yes |
| 59 | F | HBV, HCV, ASH | A | B | 15 | 144 | 145 | pos | no | yes | yes |
| 67 | M | HBV | no cirrhosis | B | n.d. | 75 | 67 | pos | no | yes | yes |
| 59 | M | ASH | A | B | 24 | 71 | 74 | pos | no | yes | yes |
| 55 | M | HCV | B | B | 26537 | 81 | n.d. | pos | no | yes | yes |
| 55 | M | HCV | A | B | 9 | 99 | 95 | pos | no | yes | yes |
| 49 | M | HCV, ASH | A | 0 | 5 | 216 | 103 | pos | no | yes | yes |
| 73 | M | HBV, HCV | A | A | 29 | 47 | 85 | pos | no | yes | yes |
| 71 | M | ASH | A | B | n.d. | 43 | 64 | neg | resection, TACE | yes | yes |
| 67 | M | NASH | A | C | n.d. | 35 | 45 | pos | resection, TACE, sorafenib | yes | yes |
| 47 | M | HCV, ASH | B | A | 3 | 31 | 34 | neg | resection | yes | no |
| 62 | M | ASH | C | B | n.d. | n.d. | 168 | pos | TACE, SIRT | yes | no |
| 52 | F | HCV | A | B | 9694 | 32 | 32 | pos | TACE | yes | no |
| 64 | M | HCV | A | A | 5 | 60 | 65 | neg | resection, 3xTACE | yes | no |
| 54 | M | HCV | A | B | n.d. | n.d. | n.d. | pos | 3xTACE | yes | no |
| 79 | M | ASH | A | B | 3 | 40 | 41 | neg | TACE | yes | no |
| 78 | M | ASH | A | C | 2 | 30 | 52 | pos | TACE | yes | no |

| Age [years] | Sex | Etiology | Child score | BCLC score | AFP [ng/ml] | GPT [U/l] | GOT [U/l] | HCMV sero-status | Treatment prior inclusion | Pheno-typic analysis | Funct-ional analysis | Patient ID |
|---|---|---|---|---|---|---|---|---|---|---|---|---|
| 67 | M | HCV | A | A | n.d. | 101 | 97 | pos | no | yes | no | HCC#1 |
| 54 | F | NASH | no cirrhoisis | B | 4 | 62 | 47 | neg | resection | yes | no | HCC#2 |
| 84 | F | NASH | A | A | n.d. | 52 | 56 | neg | no | yes | no | HCC#3 |
| 80 | M | NASH | A or no cirrhosis | B | n.d. | 64 | 46 | neg | no | yes | no | HCC#4 |
| 62 | M | ASH | A | C | 1051 | 15 | n.d. | pos | resection | yes | no | HCC#5 |
| 77 | M | ASH | A | B | 62 | 73 | n.d. | neg | no | yes | no | HCC#6 |
| 62 | M | n.d. | B | A | 13 | 28 | 59 | neg | no | yes | no | HCC#7 |
| 81 | F | NASH | no cirrhosis | A | 6 | 32 | 34 | neg | no | yes | no | HCC#8 |
| 79 | M | ASH, NASH | A | B | 71 | n.d. | n.d. | neg | 3xTACE, SBRT | yes | no | HCC#9 |
| 69 | M | HBV | A | B | n.d. | 46 | 57 | neg | 7xTACE | yes | no | HCC#10 |
| 74 | M | HBV | B | C | n.d. | 90 | 101 | neg | resection, 4xTACE | yes | no | HCC#11 |
| 70 | M | ASH | A | A | 1616 | 50 | 87 | pos | no | yes | no | HCC#12 |
| 60 | M | ASH | B | A | 10 | n.d. | n.d. | pos | no | yes | no | HCC#13 |
| 58 | M | HCV | B | A | 11 | 34 | n.d. | neg | TACE | yes | no | HCC#14 |
| 53 | M | ASH, AIH, ASH | C | A | 466 | 73 | 122 | neg | no | yes | no | HCC#15 |
| 65 | F | ASH | A | A | 5 | 14 | 20 | neg | TACE | yes | no | HCC#16 |
| 61 | M | NASH | C | B | 15 | 51 | 99 | pos | TACE | yes | no | HCC#17 |
| 55 | M | HBV/HDV | B | B | n.d. | 67 | 112 | pos | no | yes | no | HCC#18 |
| 53 | M | NASH | A | A | 3 | 46 | 32 | neg | TACE+RFTA | yes | no | |
| 80 | M | HBV | B | B | >60500 | 58 | 95 | pos | no | yes | no | |

| Patient ID | Sex | Etiology | Child score | BCLC score | AFP [ng/ml] | GPT [U/l] | GOT [U/l] | HCMV sero-status | Treatment prior inclusion | Pheno-typic analysis | Funct-Ional analysis |
|---|---|---|---|---|---|---|---|---|---|---|---|
| HCC#19 | M | HBV/HDV | no cirrhosis | B | 2 | n.d. | n.d. | neg | TACE | yes | no |
| HCC#20 | M | ASH | A | A | 6 | 74 | 47 | pos | resection | yes | no |
| HCC#21 | F | HBV | no cirrhosis | A | 27 | 23 | 34 | pos | resection | yes | no |
| HCC#22 | M | ASH | A | A | 3 | 65 | 79 | pos | TACE | yes | no |
| HCC#23 | M | HCV | A | A | n.d. | n.d. | n.d. | pos | 2xTACE | yes | no |
| HCC#24 | M | HCV, ASH | A | A | 3 | 65 | 79 | pos | resection | yes | no |
| HCC#25 | M | HBV | A | B | n.d. | n.d. | n.d. | pos | 4xTACE, SBRT | yes | no |
| HCC#26 | M | HFE | A | C | 98 | n.d. | n.d. | pos | 2xTACE | yes | no |
| HCC#27 | F | AIH | A | A | 422 | 40 | 69 | pos | resection | yes | no |
| HCC#28 | M | HBV, HCV | B | B | 298 | 189 | 209 | pos | no | yes | yes |
| HCC#29 | M | ASH | A | C | n.d. | n.d. | n.d. | pos | no | yes | yes |
| HCC#30 | M | n.d. | A | B | 9 | n.d. | n.d. | pos | no | yes | yes |
| HCC#31 | M | HBV, ASH | A | C | 7855 | 82 | 303 | pos | no | yes | yes |
| HCC#32 | M | HCV | A | A | 226 | 20 | n.d. | pos | no | yes | yes |
| HCC#33 | M | NASH | A | n.d. | n.d. | n.d. | n.d. | pos | no | yes * | no |
| HCC#34 | F | HCV | A | B | 3692 | 86 | 132 | pos | no | yes * | no |
| HCC#35 | F | n.d. | n.d. | A | n.d. | n.d. | n.d. | pos | no | yes * | no |
| HCC#36 | M | ASH | A | n.d. | 6 | 40 | 41 | pos | no | yes * | no |
| HCC#37 | M | ASH | A | B | 25 | 58 | 85 | pos | no | yes * | no |
| HCC#38 | M | ASH/NASH | A | A | 144 | 26 | 41 | pos | TACE | yes * | no |
| | M | steatosis hepatis | A | A | n.d. | n.d. | n.d. | pos | TACE, resection | yes * | no |

| Patient ID | Age [years] |
|---|---|
| HCC#39 | 71 |
| HCC#40 | 61 |
| HCC#41 | 64 |
| HCC#42 | 75 |
| HCC#43 | 68 |
| HCC#44 | 71 |
| HCC#45 | 67 |
| HCC#46 | 65 |
| HCC#47 | 76 |
| HCC#48 | 65 |
| HCC#49 | 81 |
| HCC#50 | 57 |
| HCC#51 | 70 |
| HCC#52 | 76 |
| HCC#53 | 65 |
| HCC#54 | 80 |
| HCC#55 | 77 |
| HCC#56 | 74 |
| HCC#57 | 55 |
| HCC#58 | 71 |
| HCC#59 | 79 |

**SI Table 2. Study cohort of healthy donors (HD).**

M: male, F: female, neg: negative, pos: positive

| Patient ID | Age [years] | Sex | CMV serostatus | Phenotypic analysis | Functinal analysis |
|---|---|---|---|---|---|
| HD#1 | 55 | M | neg | yes | yes |
| HD#2 | 33 | M | pos | yes | yes |
| HD#3 | 26 | F | pos | yes | yes |
| HD#4 | 34 | M | pos | yes | yes |
| HD#5 | 45 | F | pos | yes | yes |
| HD#6 | 54 | F | neg | yes | no |
| HD#7 | 56 | M | neg | yes | no |
| HD#8 | 65 | M | neg | yes | no |
| HD#9 | 55 | F | neg | yes | no |
| HD#10 | 63 | M | pos | yes | no |
| HD#11 | 61 | M | neg | yes | no |
| HD#12 | 59 | F | neg | yes | no |
| HD#13 | 54 | F | pos | yes | yes |
| HD#14 | 79 | F | pos | yes | yes |
| HD#15 | 85 | F | pos | yes | yes |
| HD#16 | 86 | F | pos | yes | no |
| HD#17 | 55 | M | pos | yes | no |
| HD#18 | 82 | F | pos | yes | no |
| HD#19 | 58 | M | pos | yes | no |
| HD#20 | 54 | F | pos | yes | no |
| HD#21 | 60 | M | pos | yes | no |
| HD#22 | 54 | M | pos | yes | yes |
| HD#23 | 53 | M | pos | yes | yes |
| HD#24 | 57 | M | pos | yes | no |
| HD#25 | 54 | F | pos | yes | no |
| HD#26 | 54 | F | pos | yes | no |
| HD#27 | 57 | F | neg | yes | no |
| HD#28 | 64 | F | pos | yes | no |
| HD#29 | 58 | F | neg | yes | no |
| HD#30 | 62 | F | neg | yes | no |
| HD#31 | 58 | F | neg | yes | no |
| HD#32 | 74 | F | pos | yes | no |
| HD#33 | 73 | F | neg | yes | no |
| HD#34 | 74 | M | neg | yes | no |

| | | | | | |
|---|---|---|---|---|---|
| HD#35 | 82 | M | neg | yes | no |
| HD#36 | 82 | F | pos | yes | no |

**SI Table 3. Study cohort of patients with liver cirrhosis.**

M: male, F: female, AIH: autoimmune hepatitis, ASH: alcohol induced steatohepatis, HBV: hepatitis B virus, HCV: hepatitis C virus, HFE: hemochromatosis, NASH: non-alcoholic steatohepatis, neg: negative, pos: positive, n.d.: not determined

| Patient ID | Age [years] | Sex | Etiology | Child score | GPT [U/l] | GOT [U/l] | HCMV serotatus | Pheno-typic analysis | Funct-ional analysis |
|---|---|---|---|---|---|---|---|---|---|
| cirrhosis#1 | 53 | M | ASH | A | 36 | 54 | pos | yes | yes |
| cirrhosis#2 | 45 | M | HBV, HCV | A | 31 | 25 | pos | yes | yes |
| cirrhosis#3 | 62 | M | ASH | C | 60 | 84 | pos | yes | yes |
| cirrhosis#4 | 48 | F | HCV | B | 46 | n.d. | pos | yes | yes |
| cirrhosis#5 | 49 | F | HCV, ASH | A | 50 | 20 | pos | yes | yes |
| cirrhosis#6 | 64 | F | HBV | B | 28 | 36 | pos | yes | yes |
| cirrhosis#7 | 49 | F | ASH | A | 15 | 32 | pos | yes | yes |
| cirrhosis#8 | 55 | M | AIH | A | 107 | 65 | pos | yes | no |
| cirrhosis#9 | 56 | M | HCV | A | 25 | 34 | pos | yes | no |
| cirrhosis#10 | 61 | F | ASH | A | 18 | 28 | neg | yes | no |
| cirrhosis#11 | 60 | F | NASH | B | n.d. | n.d. | pos | yes | no |
| cirrhosis#12 | 61 | F | ASH | B | 20 | 57 | neg | yes | no |
| cirrhosis#13 | 73 | F | NASH | C | 44 | 56 | pos | yes | no |
| cirrhosis#14 | 61 | F | HCV | A | 20 | 36 | pos | yes | yes |
| cirrhosis#15 | 58 | M | NASH | B | 58 | 44 | pos | yes | no |
| cirrhosis#16 | 74 | M | NASH | C | n.d. | n.d. | pos | yes | yes |
| cirrhosis#17 | 68 | M | ASH | B | 55 | 48 | neg | yes | no |
| cirrhosis#18 | 56 | M | ASH | B | 18 | n.d. | pos | yes | yes |
| cirrhosis#19 | 59 | M | HCV | A | 18 | 27 | neg | yes | no |
| cirrhosis#20 | 57 | M | NASH | C | 26 | 52 | pos | yes | no |
| cirrhosis#21 | 76 | M | HFE/ drug-toxic | B | 1432 | 667 | pos | yes | no |
| cirrhosis#22 | 56 | M | HBV | C | n.d. | n.d. | pos | yes | no |
| cirrhosis#23 | 71 | F | ASH | B | 39 | 59 | pos | yes | no |
| cirrhosis#24 | 59 | M | ASH | B | 38 | 82 | neg | yes | no |
| cirrhosis#25 | 68 | M | secondary biliary cirrhosis | C | n.d. | n.d. | pos | yes | no |
| cirrhosis#26 | 71 | M | NASH | C | 88 | 58 | pos | yes | no |
| cirrhosis#27 | 76 | F | NASH | A | 21 | 32 | n.d. | yes | no |
| cirrhosis#28 | 78 | F | HCV | B | n.d. | n.d. | neg | yes | no |

| | | | | | | | | | |
|---|---|---|---|---|---|---|---|---|---|
| cirrhosis#29 | 76 | M | ASH | A | 9 | 19 | pos | yes | no |
| cirrhosis#30 | 71 | F | ASH | A | 19 | 22 | pos | yes | no |
| cirrhosis#31 | 74 | M | alpha 1-antitrypsin deficiency | C | n.d. | n.d. | pos | yes | no |
| cirrhosis#32 | 68 | M | ASH | A | 19 | 27 | neg | yes | no |
| cirrhosis#33 | 66 | M | NASH | A | 29 | 17 | neg | yes | no |
